# Investigating egocentric tuning in hippocampal CA1 neurons using simulated data

**DOI:** 10.1101/2023.11.08.566253

**Authors:** Jordan Carpenter, Jan Sigurd Blackstad, David Tingley, Valentin A. Normand, Edvard I. Moser, May-Britt Moser, Benjamin A. Dunn

## Abstract

Navigation requires integrating sensory information with a stable schema to create a dynamic map of an animal’s position using egocentric and allocentric coordinate systems. In the hippocampus, place cells encode allocentric space, but their firing rates may also exhibit directional tuning within egocentric or allocentric reference frames. We compared experimental and simulated data to assess the prevalence of tuning to egocentric bearing (EB) among hippocampal cells in rats foraging in an open field. Using established procedures, we confirmed egocentric modulation of place cell activity in recorded data; however, simulated data revealed a high false positive rate. When we accounted for false positives by comparing with shuffled data that retain correlations between the animal’s direction and position, only a very low number of hippocampal neurons appeared modulated by EB. Our study highlights biases affecting false positive rates and provides insights into the challenges of identifying egocentric modulation in hippocampal neurons.

## Introduction

To orient themselves and engage in route planning, navigators must be capable of integrating incoming sensory information with their current position and combining this knowledge with existing internal representations. Thus, animals are thought to depend on the information that relates the animal’s orientation relative to points in the environment, i.e., egocentric information, and knowledge about the relationships between landmarks or distal cues, i.e., allocentric information. Information from these categories can be entirely complementary but is relayed as points on two distinct coordinate systems, which can be understood as egocentric and allocentric frames of reference. Each frame is defined by fixing the origin and rotation of the coordinate system to a particular spatial entity, such as the viewer’s head (egocentric frame of reference) or a distant landmark such as the North star or distant home location (allocentric frame of reference).^1^

Extensive experimental evidence supports the hypothesis that the hippocampus plays a crucial role in developing cognitive maps or allocentric representations of the environment.^2,3,4^ This hypothesis is primarily backed by the identification of place cells, which are hippocampal principal cells that selectively fire when an animal traverses a specific location in the environment.^2^ Place cell activity is thought to be influenced by salient distal (allocentric) cues, demonstrated by a lack of directional tuning in free-foraging tasks^5^ and field rotation in response to distal sensory cues.^6,7^ Place cells also exhibit directional selectivity when a rat navigates a maze^8,9,10^ or moves along a linear track.^11,12^

Subsequent research has identified additional cell types instrumental in the brain’s allocentric spatial representation.^13,14,15,16,17,18^ While these studies have largely focused on allocentrically tuned cells, emerging evidence indicates that many cortical regions, such as the posterior parietal cortex, lateral entorhinal cortex, postrhinal cortex, and retrosplenial cortex, represent location in an egocentric manner with reference to environmental cues, environment center, and boundaries.^19,20,21,22,23^ Recent studies have even demonstrated that the hippocampus can concurrently maintain allocentric and egocentric information during goal-directed tasks and open field foraging.^24,25,26^ This dual coding scheme in the hippocampus is somewhat unexpected, considering the extensive evidence pointing to allocentric place cells as the primary neural substrate of the hippocampal cognitive map.^2^ Considering the fact that, in open arenas, directional tuning may emerge in measures of place cell activity as a result of uneven sampling across the neuron’s place field,^9^ we decided to explore the extent to which sampling issues might account for apparent egocentric tuning in hippocampal neurons.

### Classification of directionally modulated neurons in CA1

To investigate the size of the egocentric neuronal population within the CA1 hippocampal subfield, we conducted experiments with five Long Evans rats, recording neural activity from 1,002 total cells, as the subjects foraged for cookie crumbs in an open field arena (Figure S1). The experiments were carried out in a 150×150×50 cm black box, surrounded by dark blue curtains, with a white cue card on the west wall. Of the recorded cells, 87% were spatially informative (Figure S1-S2), and 836 neurons (84%) met pre-processing criteria for inclusion in subsequent analyses.

For each neuron, we employed a classification heuristic from a study examining directionally modulated hippocampal cells (Reference-Heading (RH) model^25^). Raw directional tuning curves were fitted parametrically to a smooth cosine function to address under-sampling, effectively interpolating missing bins.^25^ This approach allowed us to determine the quantity of EB-modulated neurons in our dataset and the optimal reference points for each.^25^

We estimated egocentric bearing from either the rat’s head direction or movement direction (Figure 1A). Egocentric bearing can be described as the angular relationship between an animal’s head direction or movement trajectory and a reference point within or around its environment, which can be an object, reward, or an arbitrary location. As illustrated in Figure 1B, the firing rates of a hypothetical cell with a preferred egocentric bearing of 90 degrees show variations based on the movement’s relative direction to the reference point. In a brain region with a large fraction of spatially modulated cells such as the hippocampus, egocentric bearing must be considered in a three-dimensional sampling space, encompassing both positional and head direction coordinates (Figure 1C). While simulated behavioral data shows a high degree of coverage within this large sample space, real data exposes systematic biases, notably near boundaries.

**Figure 1:**
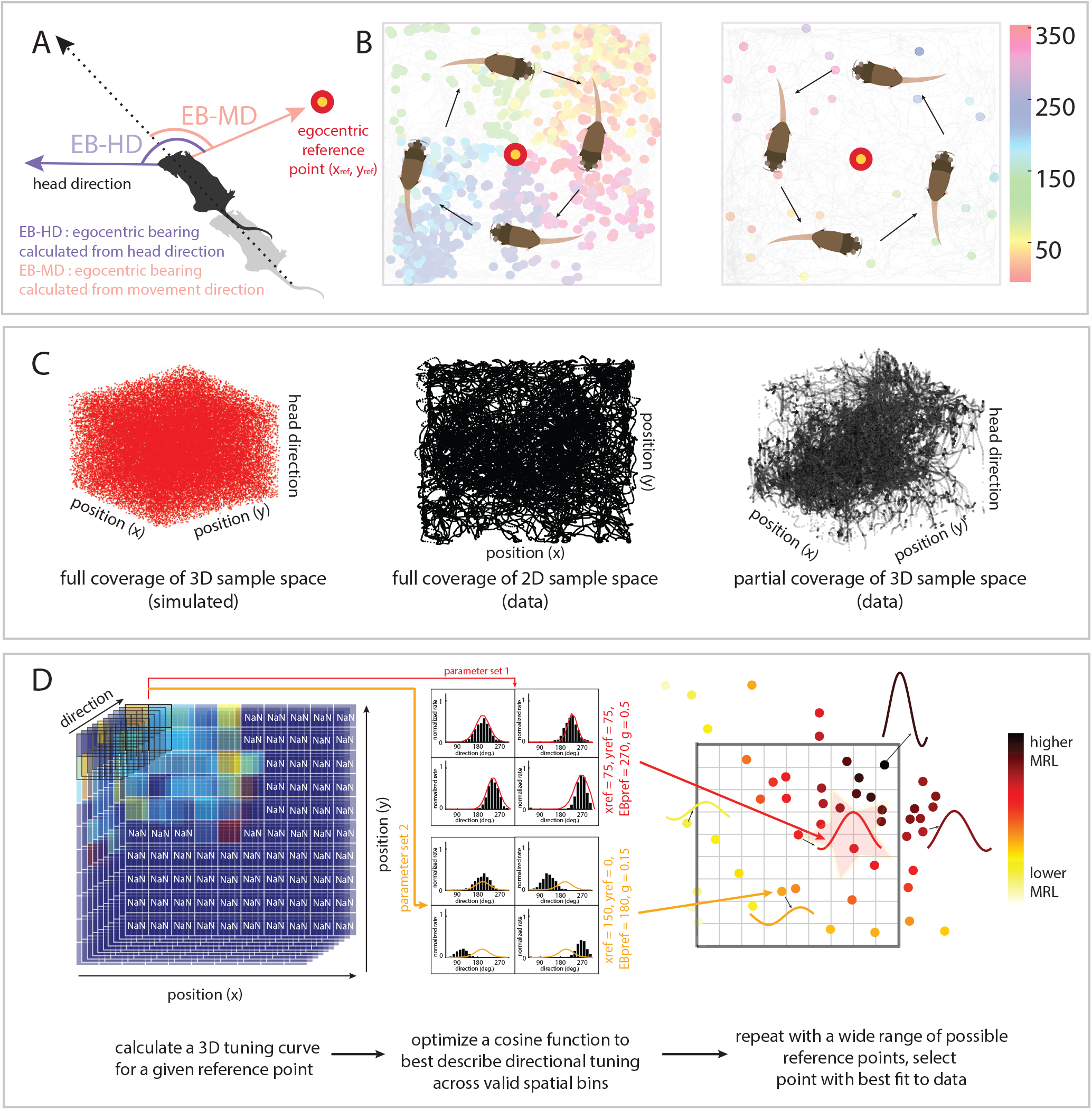
Experimental details and model of egocentric bearing. **A**. Depiction of head direction, movement direction, egocentric bearing derived from head direction (*EB*_*HD*_), and egocentric bearing derived from movement direction (*EB*_*MD*_). For each neuron *a*, Egocentric bearing (EB) is defined as the angle between the animal’s head or movement direction and a line extending from the animal’s head to a reference point (red and yellow dot) defined by coordinates (*x*_*a*_, *y*_*a*_), which may represent an object or empty location within or outside the open field. **B**. Illustration of a head-direction colored spike plot for a simulated egocentric bearing cell with a preferred egocentric bearing of 90°. Each dot indicates the cell’s spike location, with egocentric bearing at spike time color-coded by the angular variable (e.g., head direction). High cell firing rates correspond to counterclockwise movements relative to the reference point (left), whereas clockwise movements result in minimal firing (right). **C**. Egocentric bearing exists in a 3-dimensional sample space (Position_*X*_ ×Position_*Y*_ ×Head Direction). Left panel demonstrates a simulated case with a sufficiently filled sample space. In contrast, the middle and right panels present the 2-dimensional position data of a rat during a 25-minute behavioral session in a 150×150 cm open field arena, and its expansion into 3-dimensional space, respectively. Note the sampling bias, particularly near borders. **D**. The figure illustrates how the Reference-Heading (RH) model^25^ is fit to a three-dimensional tuning curve, depicted as a 10×10×10 matrix. The physical environment under consideration is discretized into 100 two-dimensional spatial bins (10×10) with each bin associated with a distinct local head or movement direction tuning curve, culminating in the construction of a two-dimensional angular tuning curve for each spatial sector in the tertiary dimension of the 3D matrix (left-most panel). Each bin value represents a normalized firing rate (Hz), denoted by a color spectrum ranging from blue (low rates) to red (high rates). The RH model aims to find the best parameters to describe the observed data using a cosine function applied to the tuning curve matrix. The iterative process of model fitting to local directional tuning curves is exemplified in the central panel, through two rounds of optimization with parameter sets #1 (red) and #2 (orange) (middle panel). The process continues until the optimal parameter set is found, which is highlighted as a set of reference points shown in the right-most panel. The optimal set minimizes the error between the model’s proposed cosine function and the cell’s empirical, discretized tuning curve, represented here by parameter set #1 (bright red tuning curve with red star around it). More information about RH model details can be found in the STAR Methods.

To estimate a neuron’s preferred egocentric bearing and corresponding environmental reference point while considering these biases, we implemented a model that utilizes 100 head direction or movement direction tuning curves, each constructed from positional data in two-dimensional spatial bins (15^2^ cm each; Figure 1D). An optimization procedure was applied to stretch, squeeze, and shift standard cosine functions to best approximate the shape of the directional tuning curve in each bin that passed firing rate and occupancy criteria (Figure 1E; STAR Methods). Due to the model’s sensitivity to initial conditions, we repeated the fitting process 100 times with varying initial conditions and selected the model with the lowest overall error.

Following estimation of the neuron’s preferred reference point (*x*_*a*_,*y*_*a*_) and egocentric bearing (θ_*a*_), we computed a metric of the strength of the tuning curve, the mean resultant length, averaged across spatial bins meeting occupancy and firing rate criteria (MRL_*a*_). This metric ranges from 0 (flatter cosine curves with less predicted directional modulation) to 1 (sharper tuning). We compared this metric with a null distribution generated by maintaining animal’s spatial path and the neuron’s spike train constant and performing one of three procedures: (1) randomly reassigning head direction indices, (2) circularly shifting the head direction estimates in time with a minimum lag of 30 seconds, or (3) utilizing head direction estimates from a behavioral session where the neuron was not recorded. We assessed whether the average mean resultant length exceeded the 95th percentile of each null distribution, indicating significant EB-modulation by the estimated parameters.

### Exploring sources of variation in counts of EB-modulated neurons

We estimated the percentage of CA1 cells modulated by ego-centric bearing (EB) under six conditions, where we employed a combination of the three permutation tests and tested ego-centric bearing either derived from the animal’s head direction (EB-HD) or movement direction (EB-MD). The choice of permutation test and directional variable significantly influenced the percentage of identified EB-modulated neurons, highlighting the importance of method selection in assessing EB-modulation in the hippocampus.

When employing a random permutation test with head direction-derived egocentric bearing (HD-RAND), a striking 89% of neurons displayed EB-modulation. However, this proportion declined to 65% and 56% upon using circular and session permutation tests,^27^ respectively (Figure 2A, first three bars). In the context of movement direction-derived egocentric modulation, the random permutation test similarly identified a high percentage of significant units (86%). Yet, the count of EB-modulated neurons markedly dropped when applying more conservative tests: 43% for circular permutation and 40% for session permutation (Figure 2A, last three bars).

**Figure 2:**
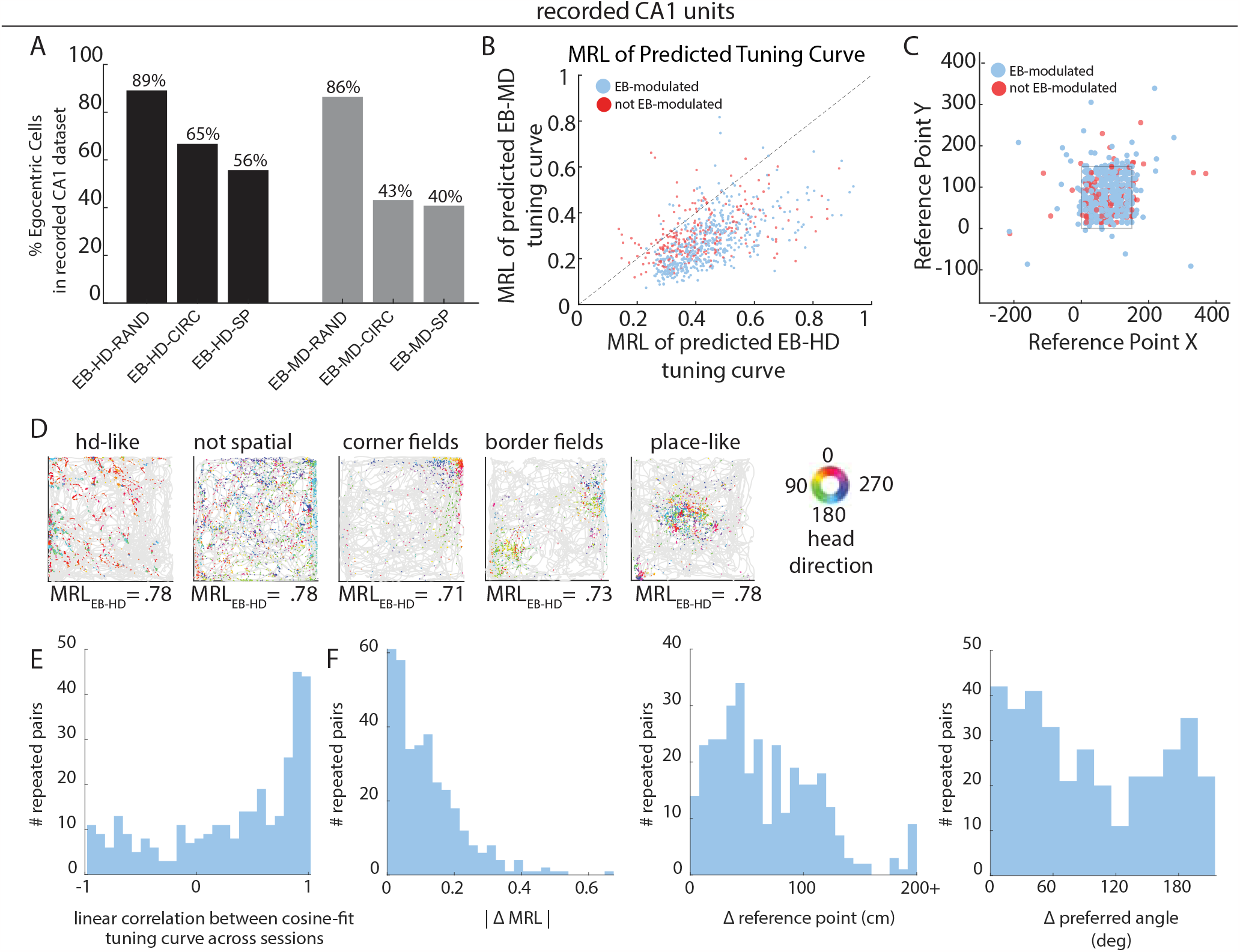
Classification of egocentric tuning in recorded hippocampal neurons. **A.** Percentage of egocentric bearing (EB) modulated cells in the recorded CA1 dataset for 62 tested combinations of angular variable and permutation test: (1) EB-HD + random (EB-HD-RAND), (2) EB-HD + circular (EB-HD-CIRC), (3) EB-HD + session (EB-HD-SP), (4) EB-MD + random (EB-MD-RAND), (5) EB-MD + circular (EB-MD-CIRC), and (6) EB-MD + session (EB-MD-SP). For example, for the first bar plot, we show the percentage of EB-modulated neurons in our CA1 dataset when egocentric bearing is calculated from head direction, and the null distribution used for hypothesis testing is built from randomly shuffling head direction indices relative to the neuron’s spike train. Results using HD are colored in black, results using movement direction are colored in grey. **B**. Mean Resultant Lengths (MRL) of RH-model predicted tuning curves for EB-HD versus EB-MD, revealing a stronger preference for EB-HD. Blue points represent significant EB-HD cells (EB-HD-CIRC), while red points are non-significant. **C**. Locations of RH model-predicted reference points for the recorded population (836 cells). The black solid line delineates recording box boundaries. **D**. Head-direction colored spike plots for six example EB-tuned units (EB-HD-CIRC), as classified by the RH-model. Grey traces display the animal’s path during a 25-min trial, while dots represent cell firing locations, color-coded by head direction at spike time. Numbers indicate tuning strength (MRL) of the RH model-fit cosine curve. Labels (e.g., ‘corner fields’) describe the spike plot phenotypes for egocentrically modulated cells. **E**. A subset of units in our CA1 dataset were recorded and tracked across multiple sessions (2+ sessions). For each pair of sessions (can be more than one pair if the cell was recorded across 3+ sessions), we calculated the linear correlation (Pearson’s r) between model estimated cosine tuning curves. **F**. Comparison of RH-model predicted parameters between session pairs for all units recorded across sessions. Left: absolute difference in predicted MRL. Middle: difference in predicted reference-heading point (x_*a*_,y*a*). Right: circular difference in predicted egocentric reference point. The wide distribution of differences suggests either limited stability of model-estimated egocentric tuning curve parameters or modulation preference remapping.

To select a single directional derivative of egocentric bearing for subsequent analyses, we compared the tuning curve strength for egocentric bearing derived from head direction (EB-HD) and movement direction (EB-MD), finding a clear preference for EB-HD (Figure 2B; p = 6.19e-150, Wilcoxon Rank-Sum Test). Consequently, we opted to define egocentric bearing as EB-HD for the remainder of our study, unless otherwise mentioned. Our analysis of all EB-modulated units revealed that reference points were consistently distributed throughout and nearby the environment, with 99% of units having a reference point < 200 cm from the center of the box (Figure 2C; distance between reference points to the center of box: median = 61.8 cm, 25th percentile = 42.4 cm, 75th percentile = 76.5 cm). This result is consistent with previous studies,^25^ where most reference points predicted by the RH model were local.

Furthermore, we discovered two unexpected properties in the detected population of neurons. First, substantial heterogeneity in firing patterns was observed across EB-modulated units with similar mean resultant length scores (Figure 2D). Second, we only found a moderate degree of stability when analyzing neurons across sessions. A subset of neurons (149 total) was tracked over 2+ sessions. For each session pair (e.g., sessions #1 and #2, or sessions #2 and #3), we calculated the linear correlation between cosine-fit egocentric tuning curves. The median Pearson correlation was *r* = 0.52, with data spanning the entire value range (Figure 2E). Only about half of the EB-modulated neurons showed a Pearson correlation over *r* = 0.8 in at least one session pair, which we took to be a strong correlation (69/149 units; Figure 2E). We also noted differences across sessions in the optimal RH model parameters: MRL (median = 0.0978, range = 0.6576), estimated reference points (median = 58.71 cm, range = 886.36 cm), and differences in preferred angles (Figure 2F), suggesting that if the cells are egocentrically tuned, tuning preferences (especially reference points) are either unstable or the tuning profiles remap across visits to an environment. Given these observations, we considered it crucial to estimate the uncertainty in the count of EB-modulated neurons.

### A simulation framework for robust hypothesis testing

To estimate the uncertainty in EB-modulated cell counts due to different choices of analysis parameters, we used a simulation approach with artificial neurons with known spatial tuning properties. By utilizing position and head direction estimates from real behavioral data, we generated spike trains for each cell using expected rates from either a two-dimensional Gaussian distribution, a von Mises distribution, or both (Figure 3A). Our simulations involved three distinct cell types thought to exist within the medial temporal lobe, namely egocentric bearing cells (EB), conjunctive egocentric bearing x place cells (EBxP), and (3) place cells (P) (Figure 3B). We also defined true positive and false positive rates, which were calculated for each experimental session, with the false positive rate (FPR) estimating the percentage of simulated place cells falsely classified as EB-modulated from a sample of 1000 units, and the true positive rate (TPR) estimating the percentage of correctly classified simulated EB or EBxP cells (Figure 3C).

**Figure 3:**
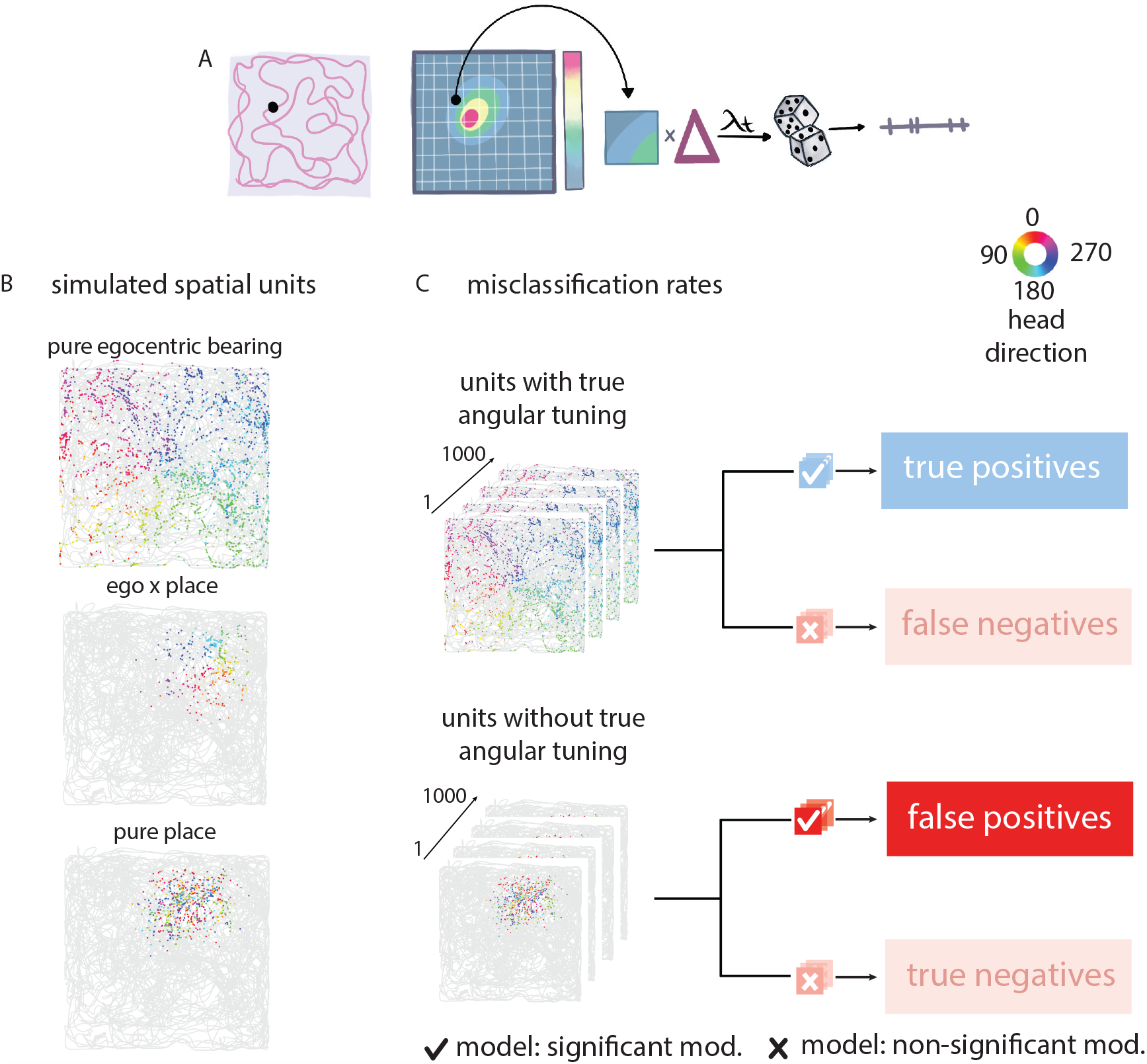
Simulation framework for validating an egocentric bearing model. **A**. Simulation procedure for functional cell types: For each timepoint (*t*_*i*_) in a recorded behavioral session, the animal’s spatial bin is determined. The average firing rate from a simulated ratemap, calculated using probability distributions (e.g., 2D Gaussian for a simulated place cell), is multiplied by an amplitude (∆). Optimal amplitude ranges are determined for each simulated cell type, and a random value is selected from this interval (STAR Methods). The product λ describes a Poisson distribution, from which a random count value is drawn and assigned to the spike train at timepoint *t*_*i*_. **B**. Head-direction colored spike plots of three simulated spatial cell types: (1) “pure” egocentric bearing, (2) conjunctive egocentric bearing x place, and (3) “pure” place. **C**. Misclassification rate calculation: Top: False negative calculation involves 1000 simulated units with true angular tuning (“pure” egocentric bearing cells) classified by the RH-model. False negative rate is defined as the percentage of units not significantly modulated by RH-angle with a local reference point. Bottom: False positive calculation includes 1000 simulated units with place tuning only (“pure” place cells) classified by the RH model. False positive rate is defined as the percentage of units significantly modulated by RH-angle.

### Variability in misclassification rates across simulated cell types

Leveraging our framework of simulated spatial units, we evaluated the true and false positive rates for each recorded open field session using each combination of directional variables and permutation tests. By generating 3000 simulated units for each session (1000 EB cells, 1000 EBxP cells, 1000 place cells), we found an exceptional performance (99.9% true positive rate) for simulated EB cells across all sessions, directional variables, and permutation tests (Figure 4A). For simulated conjunctive EBxP cells, we observed a wider distribution of true positive rates across sessions. Despite this variation, median true positive rates remained near 90% for all cases of EM-modulated neurons, indicating high but imperfect classification performance (Figure 4B). We next employed simulated place cells to estimate false positive rates for each behavioral session. We observed a broad distribution of FPRs that varied with (1) the angular variable used to calculate egocentric bearing (movement vs. head direction), (2) the permutation test applied, and (3) the session from which the behavior was recorded (Figure 4C; range of medians across conditions were 68.5%, standard deviation = 26.2%).

**Figure 4:**
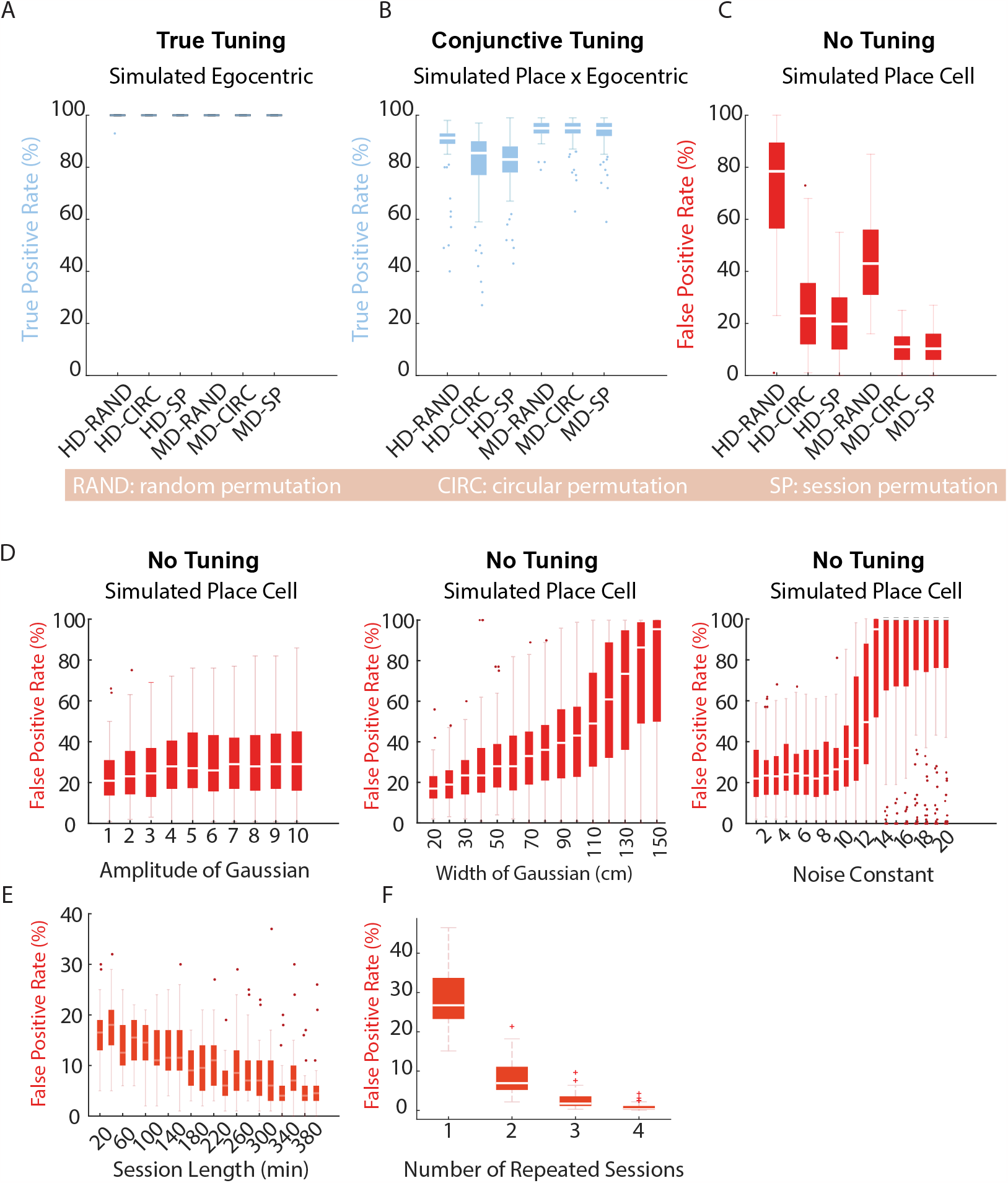
Simulated data reveals high false positive rates. **A**. Spike times for 1000 simulated egocentric bearing units are generated for each of 86 behavioral sessions (1000 x 86 total cells; open field sessions; approximatley 25 minutes each; 5 animals). True positive rates are determined for each session, with true positives defined as the number of simulated egocentric bearing cells classified as EB-modulated. Results are computed for six combinations of angular variable and permutation test applied to recorded CA1 data (see Figure 2). Each box plot data point represents the true positive rate for a single behavioral session. **B**. The same procedure as **A** was performed for conjunctive egocentric bearing x place cells to compute true positive rates. **C**. False positive rates were computed on a session-by-session basis using 1000 simulated place cells per session, with results presented as in **A** and **B. D**. False positive rates are calculated from simulated place cells as a function of increasing Gaussian amplitude (proxy for place field firing rate), Gaussian width (proxy for place field width), and Gaussian noise constant (Poisson noise). Note the significant increase in false positive rates with estimated place field width. **E**. False positive rates of simulated place cells as a function of session lengths for open field sessions. Extended sessions were made by concatenating multiple behavioral sessions. **F**. False positive rates of place cells as a function of the number of sessions a unit is recorded in. Observe that over-session stability checks reduce the false positive rate below chance (5%) for idealized place cells.

Firstly, mirroring the differences observed between EB-HD and EB-MD noted in Figure 2A, false positive rates were consistently greater when calculated using EB-HD than EB-MD (Figure 4C). These differences may be explained in part by a higher degree of mutual information between head direction and position I(P,HD) than between movement direction and position I(P,MD) (median(I(P,HD))=0.51, median(I(P,MD)=0.15; test Figure S4A-B). The Wilcoxon rank-sum test revealed a significant difference in mutual information values between the two directional variables and position (I(P,MD) and I(P,HD); W = 11.31, *p* < 0.0001). Since these mutual information measures are calculated irrespective of neural firing, it suggests that position can predict several angular variables, and the ability to predict the angular variable from position may be contributing to the disparity that we note in the false positive rates.

Secondly, we found that false positive rates varied with the permutation test applied for hypothesis testing. False positive rates were lower when hypothesis testing was conducted using circular permutation testing when compared to random permutation testing, although the median FPR across sessions was still above expected chance levels, suggesting poor performance even when using the most fitting of the field-standard permutation tests (median FPR= 23%; Figure 4C). Further, session permutation tests did not significantly improve the median false positive rate, likely due to large variations in animal behavior across sessions (W = 7.6e+03, p = 0.08, (EB-HD-SP vs. EB-HD-CIRC); W = 7.2e+03, p = 0.66, (EB-MD-SP vs. EB-MD-CIRC); Wilcoxon rank-sum test). Expectedly, FPRs were highest when random permutation testing was used for classification (median(EB-HD-RAND) = 78.5%, median(EB-MD-RAND) = 43%; Figure 3C, last three box plots). Random permutation tests are not commonly used on neural data anymore (or other data types with time-dependencies), although we noted its usage in an early publication using the RH-model which is why we included it here.^25^ We hypothesize that differences in the means and 95th percentiles of the various null distributions underlie the variation of false positive rates across permutation tests. In Figure S4C, we show a sample case where the MRL at the 95th percentile of a null distribution constructed from random shuffling of HD samples is 0.15 and is 0.27 when constructed by shifting HD samples circularly in time. Thus, because random permutation tests break time dependencies in the data, it leads to a much less conservative test.

Third and lastly, we discovered that the false positive rates fluctuate depending on the session in which the behavioral variable was recorded. Within a singular condition, such as HD-CIRC, the false positive rate can span from 0% to over 60%, with only minor variation across repeated tests (Figure S5). The classification of a simulated place cell as a false positive likely mirrors the behavioral bias inherent in the simulated units’ place field. Numerous instances of over-fitting were observed with the RH-model, often characterized by under-sampling due to consecutive missing angular bins. This under-sampling resulted in artefactual peaks in the model-fit tuning curve or cases where bimodal distributions were inaccurately fit as unimodal distributions. On a broader scale, the disparity between the model-fit cosine curve and the actual data was significantly pronounced for simulated place cells as compared to other types of simulated cells (Figure S6). Our analysis did not pinpoint any single variable with a strong linear correlation to each session’s false positive rate (Figure S4E). This aligns with expectations, as the FPR is a rate likely influenced by a combination of covarying aspects of the animal’s behavior and the simulation parameters of each unit (Figure S4F).

Given that our dataset is diverse, composed of spatial cells of all different shapes, sizes, and rates, we tested how properties of the simulated neurons influence false positive rates. We tested three parameters that govern firing rate, field size, and noise levels (amplitude, width, and noise constant of the 2D Gaussian function used to simulate place fields). We found no significant increase in the distribution of FPRs when neurons fired at low rates (median FPR = 21%; low amplitude of the simulated Gaussian) versus high rates (median FPR = 29%; high amplitude of the simulated Gaussian) (Figure 4D, first plot). However, we found significant increases in FPRs as a function of increasing place field size and out-of-field noise (field size: p=1.93e-19 smallest to the largest field width; Wilcoxon Rank-Sum Test; noise constant: p= 3.3035e-15 no noise vs. maximum noise; Wilcoxon Rank-Sum Test; Figure 4D, last two plots). The change in false positive rate with field size was the most remarkable effect, increasing from 18% for small fields to 95% when firing occupied the full extent of the 2D environment (Figure 4D, last plot). We noted that the mutual information between head direction and position I(P,HD) increased similarly when data was taken from larger portions of the environment, particularly when data from the borders was incorporated into the measure (Figure S4E). This suggests that neurons with more broad spatial tuning, and with tuning that overlaps with multiple environmental borders, are particularly susceptible to misclassification.

Given that biased sampling likely contributes to the false positive problem, we wanted to investigate how changing the experimental parameters would influence the false positive rate. To accomplish this, we examined the effect of session length and recording across repeated sessions on FPRs. Increasing session lengths from approximately 20 minutes to 1 hour reduced median FPRs to 12.5% (range at this time length = 19%; Figure 4E). We found that for average FPRs to decrease to a chance level of 5%, we had to use the maximum tested amount of data (400 minutes, 6 ½ hours) (Figure 4E). Furthermore, a cell may be unlikely to show up as a false positive in multiple, repeated sessions (Figure 4F) – suggesting that rigorous stability testing (e.g., not just testing within-session stability) should decrease false positive rates. However, in cells that do show up as FPs in 2 different behavioral sessions (when all other parameters are kept constant), the predicted egocentric reference points or other parameters may differ (see Figure 2). This issue suggests that the behavior which is sampled within the place field is capable of ‘randomizing’ where the chosen reference point is in space. With place cell remapping as an additional confound, this could present a very difficult issue for interpretation of the results.

Collectively, our findings indicate that EB-tuning is overestimated even in noiseless, simulated conditions. When such a highly flexible model (4 parameters) is used to fit data in a severely under-sampled regime, we note that the resulting tuning curves often look nothing like the data itself, often making something interesting (e.g., an egocentric tuning curve) out of noisy, NaN-filled data matrices. Further, for identification of EB-tuned cells, our findings suggest that FPRs can only be reliably mitigated by implementing extremely long, continuous recordings (approximatley 6.5 hours), which can be unfeasible for both the rodent and the researcher. A more attractive alternative would be to modify the test statistic such that the distribution against which we are testing has the same behavioral sampling issue as the data it comes from.

### Building a new kind of null distribution

We now outline an alternative approach for estimating counts of EB-modulated cells by first constructing a null distribution that considers the relationship between the animal’s position and head/movement direction, while keeping the size of the place field consistent. For each recorded CA1 cell, we generated 1000 Poisson-spiking ‘spatial copycat units’ (SPCUs) based on the neuron’s spatial firing rate map (5×5 cm bins) and the rat’s behavior during the session (Figure 5A, panels 1-2). Using the rat’s tracking data, we determined the rat’s position at each time point. We then used the neuron’s average firing rate in the spatial bin as the rate parameter (lambda) to generate Poisson spikes (Figure 5A, panel 3). The spatial firing patterns of the copycat cell are preserved as it fires at nearby, slightly varying positions from the original neuron. However, modulation of the firing rate based on head direction is not incorporated, resulting in possible inconsistencies in head direction statistics. By resampling neurons in this way, we assessed the extent of apparent egocentric direction selectivity we can expect by chance in a way that is specific to each neuron and in each behavioral session. Employing these null distributions, we calculated a cell-specific false positive rate for each neuron.

**Figure 5:**
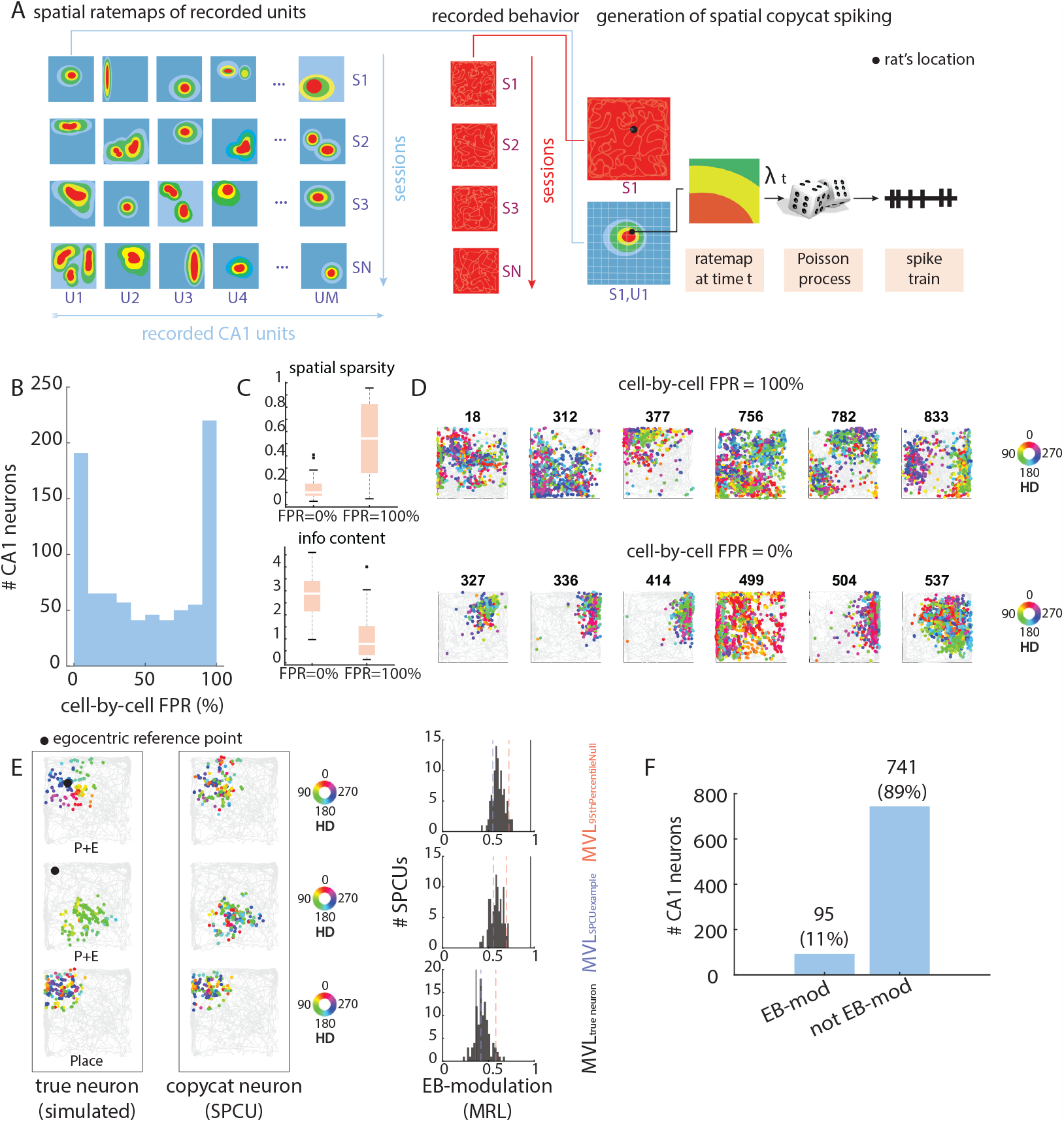
Accounting for position-direction dependencies with a null distribution of simulated ‘spatial copycat’ units. **A.** For each recorded CA1 cell, we generated 1000 simulated “spatial copycat” units by generating Poisson spiking from the recorded cell’s spatial firing rate map. By doing this, we maintained the cell’s spatial firing statistics but jumbled the head direction that the animal is in when the cell fires. By doing this, any true angular tuning should be destroyed, and we could test if the results we obtain from these simulated units are significantly different from those we compute from the true CA1 neuron. **B**. Here, we show the distribution of cell-by-cell false positive rates for each recorded hippocampal neuron. As these neurons are not endowed with any true angular tuning, this number represents the percentage of “spatial copycat” units (out of 1000 units for each recorded CA1 neuron) which are classified as EB-modulated. Note the two peaks at 0% and 100%. **C**. Spatial sparsity (top) and spatial information scores between the neuron’s spiking and the animal’s position for cells with cell specific FPRs of either exactly 0% or 100%. **D**. Head-direction colored spike plots (each colored dot represents a clustered action potential, colored by the angle of the animal’s head at the time of spiking) superimposed on the animal’s behavioral trajectory (grey) for neurons with high and low cell specific FPRs (top and bottom, respectively). **E**. This figure displays three instances of a “true” neuron, modeled either as a place cell or a combined place and egocentric cell (P+E), alongside a spatial copycat unit (SPCU) representation for each. On the left, spikes color-coded by head direction are overlaid on the animal’s gray trajectory for two P+E simulated scenarios (top two cells). A black dot indicates the egocentric reference point as determined by the RH-model. Two scenarios are presented: one with the reference point centered within the place field, generating a pinwheel effect from head-directional spike patterns, and another with the reference point outside the place field, mimicking a place cell with typical head direction tuning. Both example P+E neurons exhibit discernible directional tuning in the spike plots. The third neuron (bottom) represents a simulated place cell lacking true directional tuning; hence no reference point is provided. The second column shows examples of SPCU units, constructed using the true neuron’s spatial rate map. While overall spatial firing statistics are maintained, new spike positions are sampled according to a Poisson process. The right column showcases the distribution of mean resultant lengths (MRLs) from 100 spatial copycat cells’ model-fit tuning curves (black histograms), the true neuron’s MRL (MRL_*trueNeuron*_), the example SPCU’s MRL (MRL_*S PCUexample*_), and the 95th percentile of the null distribution (MRL95th Percentile Null). This shows that when there appears to be strong directional tuning, the MRL_*trueNeuron*_ is more likely to lie far outside of the copycat distribution. However, when there is no true directional tuning (as in the case of the simulated place cell), the test statistic (MRL_*trueNeuron*_) lies far within the null distribution. **F**. Count of CA1 neurons (out of 836 analyzed units) significantly more EB-modulated than a distribution of simulated SPCUs (tested at the 95th percentile).

We calculated the percentage of SPCUs artifactually classified as EB-modulated for each CA1 neuron, yielding SPCU-based false positive rates (SPCU-FPRs) ranging from 0-100%. The distribution exhibited two major peaks, at 0% and 100%, with a median of 49% (Figure 5A; bimodality coefficient = 0.703; skewness = 0.028; kurtosis = 1.42). The observed bimodality implies the existence of two primary cell subsets: one characterized by high confidence in EB classification outcomes, and another whose EB classification certainty may remain elusive. Only 17% of recorded CA1 cells had SPCU-FPRs below 5% (SPCU-FPR_*low*_), while 50% had SPCU-FPRs ≥50%, and 23% had *FPRs* ≥95% (SPCU-FPR_*high*_) (Figure 5B). Distributions of spatial sparsity and information content scores from the *S PCU* − *FPR*_*low*_ group were significantly lower, and significantly higher, respectively, than the group with high FPRs (median spatial sparsity: SPCU-FPR_*low*_ = 0.14, SPCU-FPR_*high*_ = 0.38; Wilcoxon Rank-Sum Test; p = 1.43e-17; median information content: SPCU-FPR_*low*_ = 2.34, SPCU-FPR_*high*_ = 1.12; Wilcoxon Rank-Sum Test; p = 1.70e-17). Consistent with our finding that FPRs (calculated at the session level) increased as a function of field size (ref. Figure 4D), we observed significantly higher spatial sparsity scores and lower information content scores for SPCU-FPR_*high*_ cells, suggesting larger spatial fields (Figure 5C-D).

Given these results, we posit that in constructing an alternative null distribution for EB classification, we should prioritize the maintenance of space-directional correlations over temporal information contained in the neuron’s spike train. We tested the null hypothesis that EB-modulated tuning does not exceed what we expect by place tuning alone given that temporal dependencies between head direction and position are maintained. EB-modulated cells were classified based on whether the EB-modulation statistic of the recorded CA1 neuron (MRL of the model-estimated tuning curve) exceeded the 95th percentile of a distribution of spatial copycat units, rather than permuting the directional variable relative to stable spiking.

In cases where there is sufficient head direction sampling within the neurons place field and the neuron has a true underlying directional tuning profile (e.g., simulated P+E neurons; Figure 5E, left column, rows 1-2), the generated SPCUs are not expected to have any discernible directional tuning (Figure 5E, middle column, rows 1-2). This is because the directional tuning is unique and intrinsic to the neuron, and not reflective of over-sampling of direction within a small area. Consequently, the MRL for true EB-modulated cells should lie far beyond the SCPU null distribution (Figure 5E; third column, rows 1-2). However, a purely spatial cell (e.g., simulated place cell) without any directional tuning should have a similarly flat directional profile to its corresponding SPCU distribution, and thus should have a MRL that lies within the SPCU null (Figure 5E; bottom row).Of 836 analyzed CA1 neurons, only 11% passed the 95th percentile SPCU criterion for EB modulation (Figure 5F).

### Mitigating false positives with adaptive significance thresholds

In a final analysis to re-assess counts of EB-modulated cells, we determined the significance levels that yield FPRs at or below 5%, This assessment was conducted at both the session level, utilizing simulated place cells, and the cell level, employing simulated spatial copycat cells. We varied the null distribution percentile that MRL must surpass to be deemed EB-modulated (100 percentiles tested at the session level, 10 tested at the cell level due to computational intensity). We then calculated FPRs for each behavioral session using simulated place cells (HD-CIRC) and selected the percentile where the FPR reached ≤5%. Only 7 out of 86 behavioral sessions had FPRs ≤5% at the commonly used 95th percentile (8% of sessions; Figure 7B). Notably, 69% (60/87) of sessions required testing at the 99th percentile or higher to achieve target FPRs (Figure 7B). For ten sessions, the FPR never reached 5%, even with the significance threshold set at the 100th percentile (Figure 7C). Eight sessions required testing at the 100th percentile.

Our initial classification (Figure 2A ‘CIRC-HD’) estimated 65% of neurons as EB-modulated. When recalculated with ‘optimal’ significance levels, only 302/836 (36%) neurons were deemed EB-modulated. We also examined the most conservative case, testing at the 100th percentile, finding 17% (142) of CA1 neurons EB-modulated (Figure 7D), slightly higher than the 11% observed when directional bias was eliminated in single-cell analyses as in Figure 5F.

We repeated these tests, calculating FPRs at the cell level, using distributions of SPCUs (1000 SPCUs per recorded CA1 neuron, 100 circular permutations per SPCU). Of 836 CA1 neurons, 134 (16%) achieved target FPRs at the default 95th percentile or lower (median=90%, mode=90%, IQR=0.0303%; Figure 7E). For 511 neurons (61%), FPRs could not be reduced below 5% (min=5.1%, max=100%, median=36.3%, mode=100%, IQR=62.1%). C-FPRs likely cannot be reliably reduced due to strong correlations between head direction and place within a cell’s place field.

Finally, we re-estimated the percentage of EB-modulated cells in our recorded dataset as a function of the significance level, testing each cell against its SPCU distribution. From the 90th to 98.85th percentile, the percentage of EB-modulated cells linearly reduced from 11% to 8% (R = −0.9904, P = 8.01e-31; 11% at the default 95th percentile). A sharp decrease to 4.5% EB-modulated cells followed when the hypothesis test was performed at the 100th percentile (Fig. 6G). Furthermore, only two of the EB-modulated cells that passed all criteria were classified as EB-modulated across successive sessions (Figure S7B). These results suggest that while egocentric tuning may be present in the hippocampus, our initial estimates, based on estimates in the published literature, vastly overstated the prevalence of EB-tuning.

**Figure 6:**
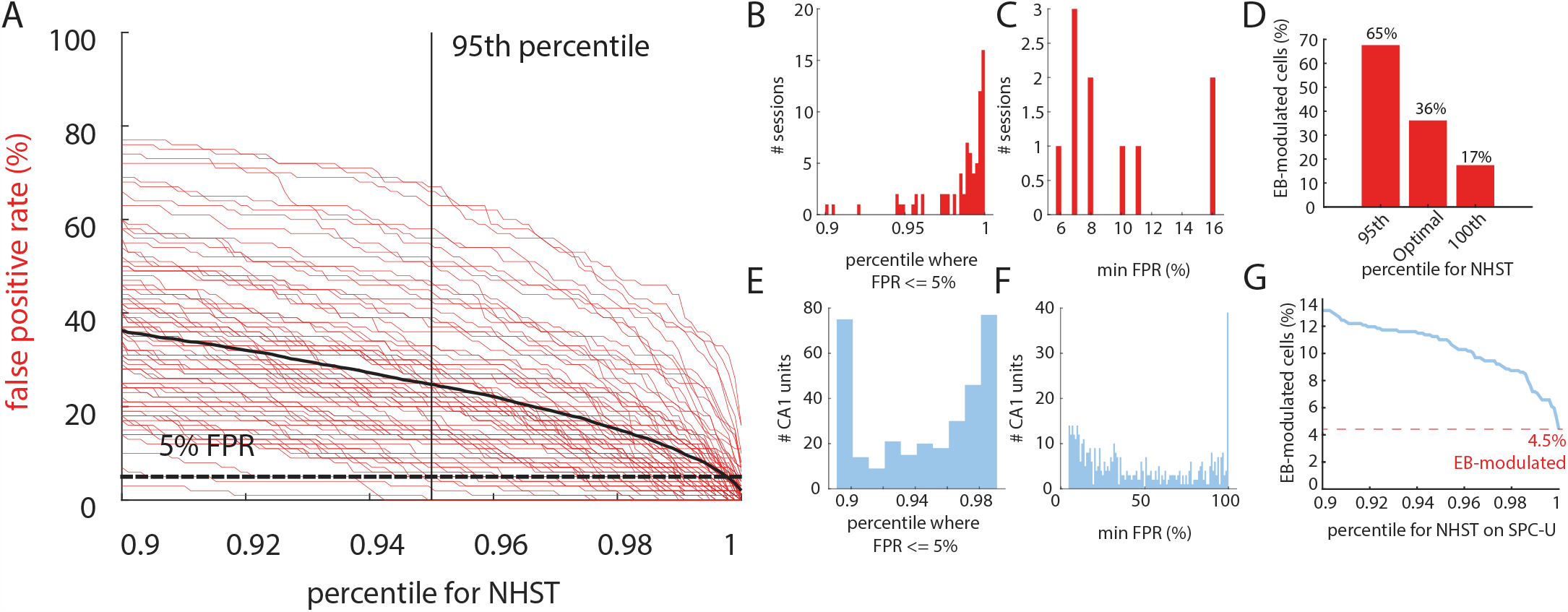
Optimizing significance levels for null hypothesis significance testing. **A**. False positive rate (FPR) as a function of percentile of the null distribution, which the true statistic (MRL of the RH-model estimated EB-tuning curve) must exceed for significance, shown for each session (grey lines) and averaged across all sessions (black line). The blue solid line indicates the FPR at the 95th percentile (our ‘default’ significance level), while the blue dashed line represents a 5% FPR. **B**. For each behavioral session, 1000 idealized place cells are generated and FPRs are calculated at 100 significance levels. Displayed is the significance level required to achieve a 5% FPR, considered acceptable, and referred to as the ‘optimal’ percentile for hypothesis testing. Ten behavioral sessions never reached a 5% FPR, even at the 100th percentile. **C**. For the ten behavioral sessions unable to achieve a 5% FPR, the minimum FPR at the 100th percentile is reported. **D**. Percentage of EB-modulated cells when tested at the 95th percentile, the ‘optimal’ percentile (derived from plots in B and C), and the 100th percentile. **E**. For each recorded CA1 neuron, 1000 ‘spatial copycat’ units (SPCUs) are generated and FPRs are calculated at ten significance levels (as in B). Shown is the distribution of the number of recorded CA1 neurons at each ‘optimal percentile’, where FPR is less than or equal to 5%. Not depicted are 511 neurons (61%) that never reached a 5% FPR, even at the 100th percentile, which are displayed in **F. F**. Same as shown in **C**, minimum FPR achieved at the 100th percentile for the 511 neurons not shown in **E. G**. For each percentile of the empirical distribution of [shuffled/null] mean resultant lengths that the true mean resultant length must surpass for statistical significance, the estimated percentage of EB-modulated cells found in recorded CA1 neurons (real data) is displayed.

## Discussion

Navigation is dependent on both world-centered and self-centered information. While world-centric information is well-studied within the hippocampal-entorhinal system3,14–18, research of egocentric directional information in CA1 neurons during free-foraging has yielded inconsistent results,^24,25,26^ leaving the true prevalence of EB-modulated cells in the hippocampus uncertain. Early studies raised concerns about the over-representation of directional tuning in hippocampal neurons without task or environmental geometry,^9^ suggesting that head direction tuning could be attributed to biased sampling of head angles within a cell’s place field. Our study aimed to quantify EB-modulated neurons while accounting for potential false positives to clarify the fraction of cells encoding egocentric spatial information, resolving discrepancies in previous literature, and highlighting factors leading to artefactual tuning detection.

We investigated egocentric directional signal encoding by hippocampal pyramidal cells during open-field foraging. Although we found support for an egocentric signal in a small subset of neurons, our results suggest that EB-modulation counts in the hippocampal CA1 region have been substantially overestimated. Our pre-correction estimates align with a similar study on reference-heading tuning in mice (56% total RH-modulated neurons; open field recordings^25^). After accounting for false positives, however, we found that at most, 17% of the population was EB-modulated (Figure 6D; estimated % of EB-modulated cells when the test statistic is tested against a null distribution of circularly shuffled HD values at the 100th percentile). While we do not directly test goal directed behaviors here, our results are consistent with egocentric tuning that has been shown to arise during reward-seeking behavior. Our results concur with findings that population-level egocentric tuning was robust in goal-directed tasks but low during open-field foraging^26^ (31% during goal task, 13% during foraging). Similarly, only 18% of CA1 cells in bats (58 out of 309) were reported to be tuned to an egocentric direction relative to a goal location.^24^ Nevertheless, when accounting for false positives at the cell level, our most conservative and seemingly reliable estimates reveal that only 4% of the recorded population in the open field was EB-modulated (Figure 6G) when true neurons were tested against 100th quantile of a distribution of simulated cells made with comparable spatial tuning profiles (SPCUs), which is within chance levels when hypothesis testing at a field-standard significance level.

Our investigation of EB-modulation estimates stems from a series of tests exhibiting considerable variation in misclassification rates. These rates are contingent upon several factors, including the directional variable defining egocentric bearing, permutation test, hypothesis testing significance level, and the parameters of simulated spike trains. Based on our analyses, we propose four primary conclusions.

Firstly, our findings emphasize that movement direction is not an appropriate proxy for head direction, despite its previous use as such.^24,25^ Secondly, the null distribution employed to test egocentric tuning in spatial neurons must adequately account for correlations between tested variables (e.g., position and directional variables).^28^ Thirdly, when using highly flexible approaches, such as the RH model or similar techniques (e.g., grid searching for an optimal reference point), it is crucial to test at a conservative significance level (e.g., does the null distribution exceed the 99th or 100th percentile?) that mitigates false positives to identify novel tuning. In this context, a highly flexible approach refers to adaptable models with several parameters (like the RH-model), which can be useful for fitting complex data spaces (e.g., 3D) and addressing challenges such as interpolating over numerous empty bins in a dataset. However, while these approaches can be very useful, they are also prone to overfitting, and should be subjected to rigorous validation when applied to new problems and datasets. Lastly, extending the length of recording sessions or conducting multiple, repeated sessions should help alleviate false positives; however, this approach does not guarantee complete elimination of uncertainty.

Our study provides a more accurate estimation of EB-modulated cells in the hippocampal CA1 region by rigorously accounting for potential false positives. We have not only addressed inconsistencies in the existing literature but also offered key insights into the factors affecting the detection of egocentric directional tuning in spatial neurons. By doing so, this study contributes to a better estimate of the actual prevalence of cells encoding self-centered spatial information within the hippocampal system.

## Acknowledgements

We thank S.O. Andersson for important discussions and manuscript comments. M. Guardamagna and V.L. Kargård Olsen for manuscript comments. K. Haugen performed histology, imaging, and anatomical discussions on tetrode locations. E. Kråkvik, I. Ulsaker-Janke, D. Lederberg, and J. Sugar performed and assisted J.S. Blackstad during surgeries. This work was supported by funding from the Research Council of Norway (FRIPRO grant number 300394 to M.B.M., Centre of Excellence grant number 223262 to M.B.M. and E.I.M., National Infrastructure grant number 295721 to E.I.M. and M.-B.M); a grant from the K.G. Jebsen Foundation (grant number SKGJMED-022); the Kavli Foundation (M.B.M. and E.I.M.); a direct contribution to M.B.M. and E.I.M. from the Ministry of Education and Research of Norway; and the Research Council of Norway - iMOD, NFR grant number 325114 (B.A.D).

## Author Contributions

J.S.B., E.M., and M.B.M. conceived experiments. J.S.B. conducted animal experiments. B.A.D. and J.C. developed analyses. J.C. analyzed the data and made figures. D.T. provided analytical advice and support through the project, and V.N. provided discussions. J.C. wrote the manuscript with advisory support from E.M., B.A.D., and M.B.M. B.A.D., E.I.M. and M.B.M supervised the project; B.A.D., E.I.M. and M.B.M. obtained funding.

## STAR Methods

### RESOURCE AVAILABILITY

#### Lead Contact

Further information and requests for resources and reagents should be directed to and will be fulfilled by Edvard Moser and May-Britt Moser. Further information about data analytics should be directed to Benjamin Dunn.

#### Data and Code Availability

The code generated during this study will be available on GitHub, and the data will be available upon request following peer-reviewed publication.

### EXPERIMENTAL MODEL AND SUBJECT DETAILS

#### Animal handling and behavioral training

Five adult male Long Evans rats, (age = 3-6 months) were used in this study. Animals were handled for several days prior to experimental training. After handling, animals were exposed to a black 150 x 150 cm recording arena (wall height = 50 cm) for several days before recording. Per the senses of the experimenter, the recording environment remained unchanged throughout the experimental period. It was characterized by blue drapes hanging around the recording box, a white cue card on the west wall and a hardware setup on the south wall. Animals were recorded in a standard ‘open field’ session in which they foraged for chocolate cereal crumbs that were sprinkled on the arena floor by the experimenter. The open field was 150×150×150 cm in size, with black walls and floors. Dark blue curtains surrounded the open field, and a white cue card was placed on the West wall of the box. The same animals were also trained on a spatial task which was not included here.

#### Animal handling and behavioral training

Before surgery, animals were housed in enriched cages with 3-8 male littermates. Following surgery, rats were moved to individual plexiglass cages (plexiglass, 45×44×30 cm) and maintained on a 12h light/12h dark schedule in a temperature and humidity-controlled facility. All experiments were performed during the dark phase. Animals were food deprived during behavioral training and weight was monitored daily to be maintained above 90% of baseline body weight. During the experiments all animals either maintained their baseline or increased weight. Water was provided ad-libitum. All experiments and surgical procedures were performed in accordance with the Norwegian Welfare Act and the European Convention for the Protection of Vertebrate animals used for Experimental and other Scientific purposes. Permit numbers 7163 and 18011.

### METHOD DETAILS

#### Surgical Procedures

Rats were anesthetized with 5% isoflurane (flow rate = 1.0 L/min, concentration = 0.75-3%) and implanted with either a 14-tetrode hyperdrive (left hemisphere, 1 animals) or two 4-tetrode microdrives (Axona LTD, one in each hemisphere, 4 animals) targeting the dorsal CA1 region of the hippocampus. Tetrodes were constructed from four twisted 17-micrometer polyimide coated platinum-iridium (90%-10%) wires (California Fine Wire). Electrode tips were plated with platinum to reduce electrode impedance (target impedance = 120-220 kΩ at 1 kHz). Tetrodes were implanted in the cortex above the target area (hyperdrives at 940 µm ventral to the dura, and Microdrives at 1500-1600 µm ventral to the dura). Implants were secured by dental cement (Meliodent), Optibond glue, and 2-8 screws (size M1.4) depending on size of the implant. Two additional stainless-steel screws in contact with cerebrospinal fluid were used for electrical grounding.

#### Behavior and Data Processing

Recordings were conducted using a Axona Ltd daqUSB system for Microdrives and Neuralynx system with Cheeta software for hyperdrive recordings. In both cases, the implanted drives were connected to the recording system using a counterweighted lightweight cable to further reduce the load on the animal’s head. An overhead camera recorded the position of two head-mounted LEDs (1 blue, 1 red) on the head stage at a sampling rate of 50 Hz (distance between diodes = 3 cm). Electrodes were gradually lowered by a maximum of 100 µm per day (average depth of CA1 recording = 2300-2600 µm, measured from the surface brain’s surface includes recordings from the time point tetrodes judged to be in the CA1 region of the hippocampus, evaluated by using local field potential markers such as the presence of the theta oscillation and sharp waves, and the presence of spatially tuned units. Unit activity was amplified by a factor of 6000 – 10 000 using either an Axona or Neuralynx preamplifier, and bandpass filtered at 600–6,000 Hz (Neuralynx) or 800–6,700 Hz (Axona). Spike waveforms within a threshold of 50-80 µV were recorded and timestamped at 32 kHz (Neuralynx) or 48 kHz (Axona). Local field potentials were amplified by a factor of 250 – 3000, low-pass filtered at 300-475 Hz and sampled at 1800-2500 Hz. Waveform extraction and initial clustering was conducted manually using MClust (A. David Redish). Following unit isolation and extraction of units with a mean spike rate of > 1 Hz and < 10 Hz across a session were classified and considered for the following analyses (836 of 1002 units). Position estimates were convolved with a Gaussian kernel (σ = 3) and a speed threshold was applied to avoid detecting brain activity associated with offline processing (such as firing related to hippocampal sharp waves) or noise activity (reflection of LEDs, movement of animal by experimenter, etc.). The animal’s running speed was calculated for each video frame and speeds below 6 cm s^−1^ or above 60 cm s^−1^ were removed.

### QUANTIFICATION AND STATISTICAL ANALYSIS

#### Estimating directional variables from animal tracking

We calculated both movement and head direction of the rats using the tracking LEDs. For movement direction, the movement direction (M) was computed from the center points (*x, y*) of the rat’s head, as *M* = atan2 (∆*y*, ∆*x*), where ∆*x* and ∆*y* denotes the changes in the head-position between consecutive video frames after mild smoothing. Head direction was calculated from the positions of the two LEDs (*x*_1_, *y*_1_) and (*x*_2_, *y*_2_) as *H* = atan2 (*y*_2_ − *y*_1_, *x*_2_ − *x*_1_).

#### Estimating tuning curves for egocentric bearing

First, we note that for a small enough spatial bin that is sufficiently far away from the preferred location of a neuron with egocentric bearing selectivity, the head direction tuning curve (i.e. using only data from when the animal was in that spatial bin) will align with the egocentric bearing tuning curve when the preferred angle and location are known. If we assume that the preferred angle and location are constant for the entire environment, one should be able to estimate those by modeling the tuning curves across multiple spatial bins. This is the intuition behind the approach developed by Jercog et al., which we describe in this section.

Let *t*_*i j*_ be the set of time bins such that *x*_*i*_ −∆_*x*_ < *x*_*t*_ <= *x*_*i*_ + ∆_*x*_ and *y* _*j*_ −∆_*y*_ < *y*_*t*_ <= *y* _*j*_ +∆_*y*_ with *i, j* = 1, 2, …, 10 and *x*_*i*_, *y* _*j*_, ∆_*x*_, ∆_*y*_ determined such that the (1.5 x 1.5 meter) environment is divided evenly into 100 equally-sized spatial bins. One can construct, for each neuron *a*, head direction (or movement direction) tuning curves which we will generalize here as 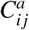, using the typical approach of binning in the angular dimension and dividing the number of spikes in each angular bin by the time spent in that bin. Here, however, 100 such angular tuning curves are constructed, one for each set of time bins *t*_*i j*_. Egocentric bearing tuning curves for each spatial bin can then be estimated by assuming a simple tuning curve shape, with four parameters: a gain parameter (*g*_*a*_), a preferred location (*x*_*a*_, *y*_*a*_), and preferred angle θ_*a*_, and minimizing the difference of those to the normalized tuning curve in each spatial bin:

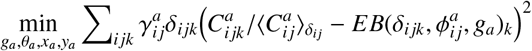

where k is the index of 10 equally-spaced angular bins, ∆_*i jk*_ is an indicator equal to 1 if the time spent in *k*th angular bin in the (*i j*)th spatial bin both exceeds a threshold of 500ms, and 0 otherwise. 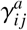 is also an indicator function that is 1 if Σ_*k*_ ∆_*i jk*_ > 5 and if the average firing rate of the neuron *a* in spatial bin (*i j*) exceeds 0.5Hz.

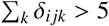is the average over angular bins with sufficient occupancy 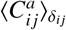,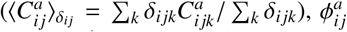 is the angular difference between the angle in allocentric coordinates from the bin center to the reference location and the preferred angle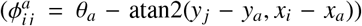 and *EB* (·) is a model of the egocentric bearing tuning curve:

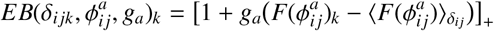

Where [·]_+_ is the Heaviside function and *F* (·) is a function chosen to constrain the shape of possible tuning curves to a cosine shape, 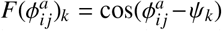, where ψ_*k*_ are the angular bin centers of the 10 angular bins. The model is fit by minimizing the squared difference between the model’s prediction and the data, summed over all bins that meet the occupancy and firing rate criteria.

The optimization was performed using MATLAB’s *fminsearch* function, which follows a Nelder-Mead simplex algorithm approach to minimize the error function. Since this model is sensitive to initial conditions and often converges to local minima, the optimization was performed 100 times for each unit, with varying initial conditions. The initial conditions are set by sampling *g*_*a*_ uniformly on the interval (0, 1), θ_*a*_ as an integer on [−180, 180) sampled uniformly and *x*_*a*_, *y*_*a*_ randomly from the set of spatial bins centers.

The iteration that yielded the lowest overall error was taken as the global minima, and the best-fit values for the four parameters were selected accordingly.

We note that if the data suggests a flat tuning curve, normalizing it by the mean (as above) would give it a constant value of 1 which would then correspond to an estimated *g*_*a*_ of 0. The normalization in each spatial bin also should account for spatial selectivity confounds, such as in the case of place cells. Furthermore, the simple shape of the cosine tuning curve and the overall low number of parameters keep this seemingly problematic optimization problem reasonably well-behaved and efficient.

#### Determining significance for results of RH model

We assessed the significance of the model’s predictions, thereby classifying a given neuron, *a*, as either (1) significantly EB-modulated or (2) not significantly EB-modulated. We calculate a metric of the model-estimated egocentric bearing (EB) tuning curve by using the mean resultant length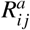 for each of the EB-tuning curves for each spatial bin that passes the occupancy and firing rate threshold criteria *EB*_*i*_. The mean resultant length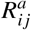 for each EB tuning curve is calculated as follows:

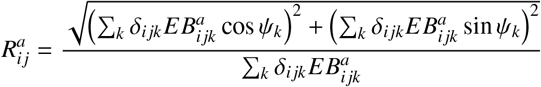

where 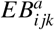 is the *k*th bin of the egocentric bearing tuning curve for spatial bin (*i j*). Subsequently, the mean resultant length for all tuning curves, now denoted as *MRL*, is computed as:

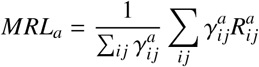

This *MRL*_*a*_ score represents the average mean resultant length of the EB tuning curves across all spatial bins and is used as the test statistic to be compared with a null distribution.

To examine the impact of various permutation tests, we keep neural spike times static and generate a null distribution by either (1) randomly reallocating head direction or movement direction indices (random permutation test), (2) circularly rotating the head/movement direction time series in relation to neural spike times (circular permutation test) with a minimum shift of 30 seconds, or (3) applying head/movement direction from a different recorded session (session permutation test). To create a single null unit for the circular permutation test, the function *h*[0 : *T*] is substituted with *h*[*s* : *T* ; *h*_0_ : *s*] where *s* is an element of *E*, ≤300 *s* ≤ 3000. If *MRL* surpasses the 95th percentile (determined in a non-parametric manner) of the null distribution of *MRL* values, the results are regarded as statistically significant. Further, if the model’s predicted reference point is < 200 cm from the center of the recording arena, the cell is taken to be egocentric, and if the reference point is > 200 cm from the center of the recording arena (a more distant reference point), it is taken to be tuned to head direction.

#### Neurons over sessions

As recording neurons across sessions was not the primary focus of this study, we found a subset of 68 neurons that were tracked in one or more open field session. The neurons were determined to be the same by manual waveform and cluster position matching, and correlating the place field locations across sessions, which typically remain consistent across multiple exposures to the same environment.

#### A case of spatial, but not directional, tuning

We independently generate large sets of simulated place cells based solely on spatial constraints to simulate the case that a neuron has spatial tuning but does not encode any directional information (e.g., head direction or egocentric bearing). We use position and head direction estimates from recorded behavioral sessions to accomplish this. We assume that each neuron spikes according to an inhomogeneous Poisson process with firing rate determined by a simulated spatial tuning curve, evaluated at the instantaneous position of the animal.

Using binned position information over time, where each bin corresponds to a location in the recording arena, and a set of user-defined parameters, spatial rate maps *G*(*x, y*) were generated for faux place cells using a Gaussian probability distribution function such that

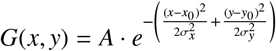

where the probability of observing a spike decreases as the animal moves farther from the center of the place field. The user-defined parameters are as follows: (*x*_0_, *y*_0_) is the center of the “place field” where *x*_0_*andy*_0_ ∼ U (3, 17). Values near borders (outer 10% of arena) are excluded due to large behavioral biases in those areas. The spread of the place field is defined by (σ_*x*_, σ_*y*_), where σ_*x*_, σ_*y*_ ∼ U (12, 18). Finally, we define an amplitude scaling factor, *A*∼ U (6, 8). The generated spike train is then given by *S* _*t*_ ∼Poisson(λ_*t*_), where λ_*t*_ = *G*(*x*_*t*_, *y*_*t*_)∆*t*, and ∆*t* is the time per video frame (0.20 s). In cases where noise is present in simulations, a constant, ϵ, is added to λ_*t*_, such that *S* _*t*_ ∼ Poisson(λ_*t*_ + ϵ).

#### Simulating additional spatial cell types

Two additional populations of spatially informative cell types were generated to test the specificity of the applied classification heuristics: egocentric bearing cells and conjunctive place cells. To generate spiking for a putative egocentric bearing cell, a reference point (*x*_ref_, *y*_ref_), a preferred egocentric orientation (θ, ranging from 0 to 360°), and peak amplitude *A*, in arbitrary units) was randomly generated. Possible amplitude values were bounded between 5 and 20 and reference points were restricted such that only values that fell within 90% of the internal segment of the arena. The shape of the von-Mises probability distribution (concentration value, κ∼ U (5, 10)) was then used to model the response curve of the neuron:

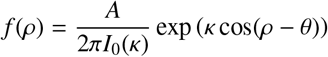

where A is a scalar, *I*_0_ is a zero-order modified Bessel function first of kind and ρ = atan2(*y*− *y*_*a*_, *x*− *x*_*a*_) −HD. For each time point, the animal’s egocentric orientation relative to the reference point θ and a corresponding spike probability, *f* (ψ), was determined. A spike count was then randomly generated from a Poisson distribution specified by the rate parameter at each time point *t*, λ_*t*_ = *f* (ρ_*t*_)∆*t*.

Conjunctive place× egocentric cell (P× E) populations were modeled by multiplying together the two components, λ = *G*(*x, y*) × *f* (ρ). Thus, the rate parameter of the Poisson distribution from which a spike count is randomly drawn is determined by the joint probability distribution of the spatial and egocentric bearing variables.

#### Hypothesis testing using simulated units

To test the null hypothesis that there is no relationship between a cell’s spiking activity and egocentric bearing on some coptimal reference point, we followed a typical null hypothesis significance testing (NHST) procedure.

A large set of place cells were simulated using the position and head/movement direction behavior from 86 open field behavioral sessions (1000 units/session). The session-by-session false positive rate was defined as the percentage of simulated place cells (which are generated with no true directional tuning) for which the results from the RH-model were considered statistically significant according to the procedure described in previous sections (unit is “EB-modulated”).

True positive rates were defined as the percentage of simulated egocentric bearing cells (initialized with true directional tuning) for which the results from the RH-model were correctly considered statistically significant.

#### Calculating mutual information between direction and position

Increasing amounts of head direction, movement direction, and position data were taken from concentric circular bins (radii = 1-100 cm) emanating from the center of the environment. With this, down-sampled data, position, and direction (either movement direction or head direction) were binned (900 bins of 5×5 cm for position, 10 bins for head direction). The mutual information between the two variables *I*(*X*; *Y*), where *X* is position and *Y* is either head or movement direction was calculated as follows:

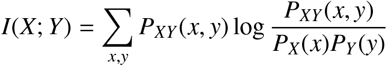

Where *P*_*X*_(*x*) and *P*_*Y*_ (*y*) are the marginal probabilities *P*_*X*_(*x*) =Σ _*y*_ *P*_*XY*_ (*x, y*) and *P*_*Y*_ (*y*) =Σ _*x*_ *P*_*XY*_ (*x, y*), and *P*_*XY*_ (*x, y*) is the joint probability distribution.

#### The intuitive approach: cell-wise false positive rates

We create a pseudo-null distribution for each recorded CA1 neuron based on the cell’s spatial firing rate maps. By comparing the directional tuning in the true case to that observed in a large set of ‘spatial copycats’, we can get some intuition about the robustness of a neuron’s directional tuning. In the case where directional tuning is *not confounded* by behavior (and is truly robust), we should effectively destroy (or greatly reduce the strength of) directional tuning in the pseudo-null distribution since the simulation parameters do not depend on head direction (or egocentric bearing, or movement direction) in any way. Thus, for each recorded CA1 neuron, we create a set of 1000 simulated spatial copycats, carry out the classification heuristics noted in the previous methods sections, and compare the resulting *MVL*_*true*_ with the distribution of *MVL*_*copy*_.

#### Session length analysis

Behavior was concatenated across multiple open field sessions to simulate data for session length analyses. A desired session length was determined, and session indices (1 through 86) were pulled from a uniform distribution of integers.

#### Repeated Sessions Analysis

To investigate the influence of recording cells across varied behavioral sessions on false positive rates, parameters defining the field width, firing amplitude, and place cell center were established for 1,000 simulated neurons. These parameters were systematically applied to reproduce the properties of the neurons using the animal’s position behavior from 86 open field sessions. Then, the RF-model was applied to the 1,000 simulated neurons across all sessions. Using this data, we first calculated the percentage of behavioral sessions in which any given neuron (e.g., simulated neuron #1) was categorized as a false positive, providing the false positive rate for when the Number of Repeated Sessions equals 1. Following this, the formula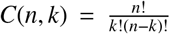 was used to compute all potential combinations of *k* sessions from the total of *n* sessions. This calculation allowed us to examine the simulated cell’s behavior across every plausible combination of 2, 3, or 4 behavioral sessions from the total 86. For each simulated neuron, its classification as a false positive was assessed in each session combination, yielding the false positive rates when the Number of Repeated Sessions is greater than 1.

#### Correlations between behavior and false positive rates

For each experimental session, we calculated several parameters, including position, head direction (HD), angular velocity, speed, acceleration, and their various statistics (e.g., mean, median, standard deviation). Circular statistics were applied to the angular data to obtain metrics such as mean, median, and standard deviation. We also calculated variance, skewness, and kurtosis for angular velocity, speed, and acceleration. Following this, we calculated the Pearson r correlation between the various statistics and the false positive rate for each given session. To investigate how the various statistics of behavior co-varied, we constructed cross-correlation matrices, which takes its absolute value to capture the magnitude of correlation irrespective of direction. After calculating this correlation matrix, hierarchical clustering is applied on the distance metrics derived from the correlations using the average linkage. The correlation matrix is reordered based on the optimal leaf ordering, which minimizes the distance between closely correlated variables.

#### Statistical Tests

We used Wilcoxon rank-sum tests for paired tests. Correlations were determined using Pearson’s *r* correlation coefficients. No statistical methods were used to pre-determine sample sizes. The experimental study contained no randomization to experimental treatments and no blinding.

---

**Table 1:** KEY RESOURCES TABLE.

| Resource | Source | Identifier/Location | Description |
| --- | --- | --- | --- |
| **SUBJECTS**<br>Rat Long-Evans | Charles River | https://www.criver.com/ | 5 male adults |
| **DATA AND SOFTWARE**<br>Simulation toolbox | Jordan Carpenter | https://github.com | Available after publication |
| CircStats | Philipp Berens | mathworks.com/matlabcentral/fileexchange | Circular statistics |
| CA1 dataset | Jan Sigurd Blackstad | N/A |  |
| MClust | A David Redish | github.com/adredish | Cluster cutting |
| MATLAB | MathWorks | https://www.mathworks.com/ |  |
| NumPy | NumPy | https://numpy.org/ | Data processing |
| SciPy | SciPy | https://scipy.org/ | Data analysis |
| Seaborn/Matplotlib | seaborn/matplotlib | https://seaborn.pydata.org/ | Data visualization |
| **OTHER**<br>Axona recording system | Axona Ltd. | http://www.axona.com/ | Hyperdrive animals |
| Neuralynx data acquisition system | Neuralynx | https://neuralynx.com/ | Microdrive animals |
| 17-micrometer polyimide coated platinum-iridium wires | California Fine Wire | https://calfinewire.com/ | Tetrode construction |
| Acquisition Software (Neuralynx) | Cheetah | https://neuralynx.com/research-software/cheetah/ | Electrophysiology Recording & Experiment Control |

## Supplemental Materials

**Supplemental Figure 1:**
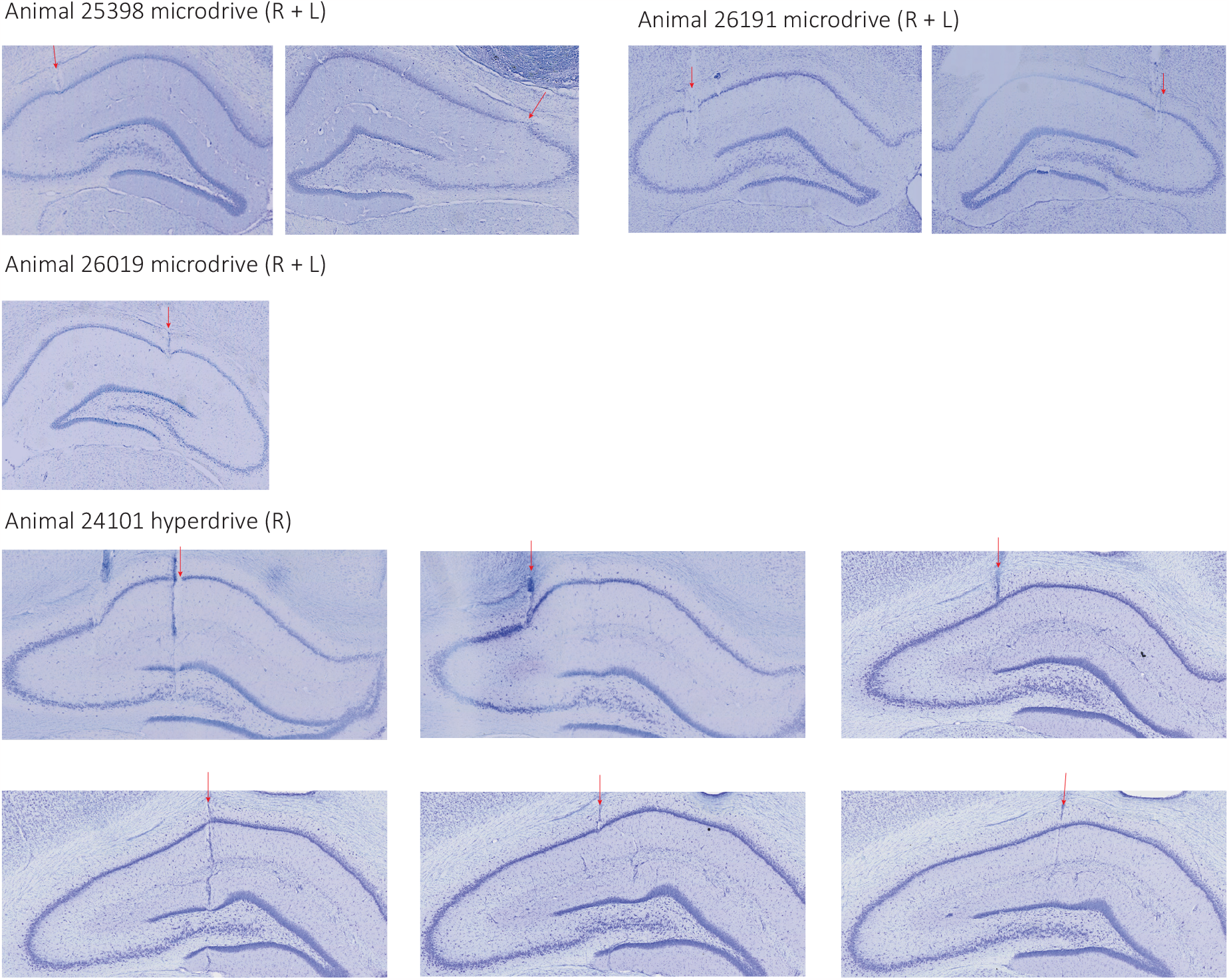
Histology and Experimental Setup. **A**. Coronal Nissl-stained sections showing recording locations in the hippocampus. The red arrow indicates the observed tetrode tract. The black and yellow dashed lines delineate the extent of the CA1 subfield of the hippocampus.

**Supplemental Figure 2:**
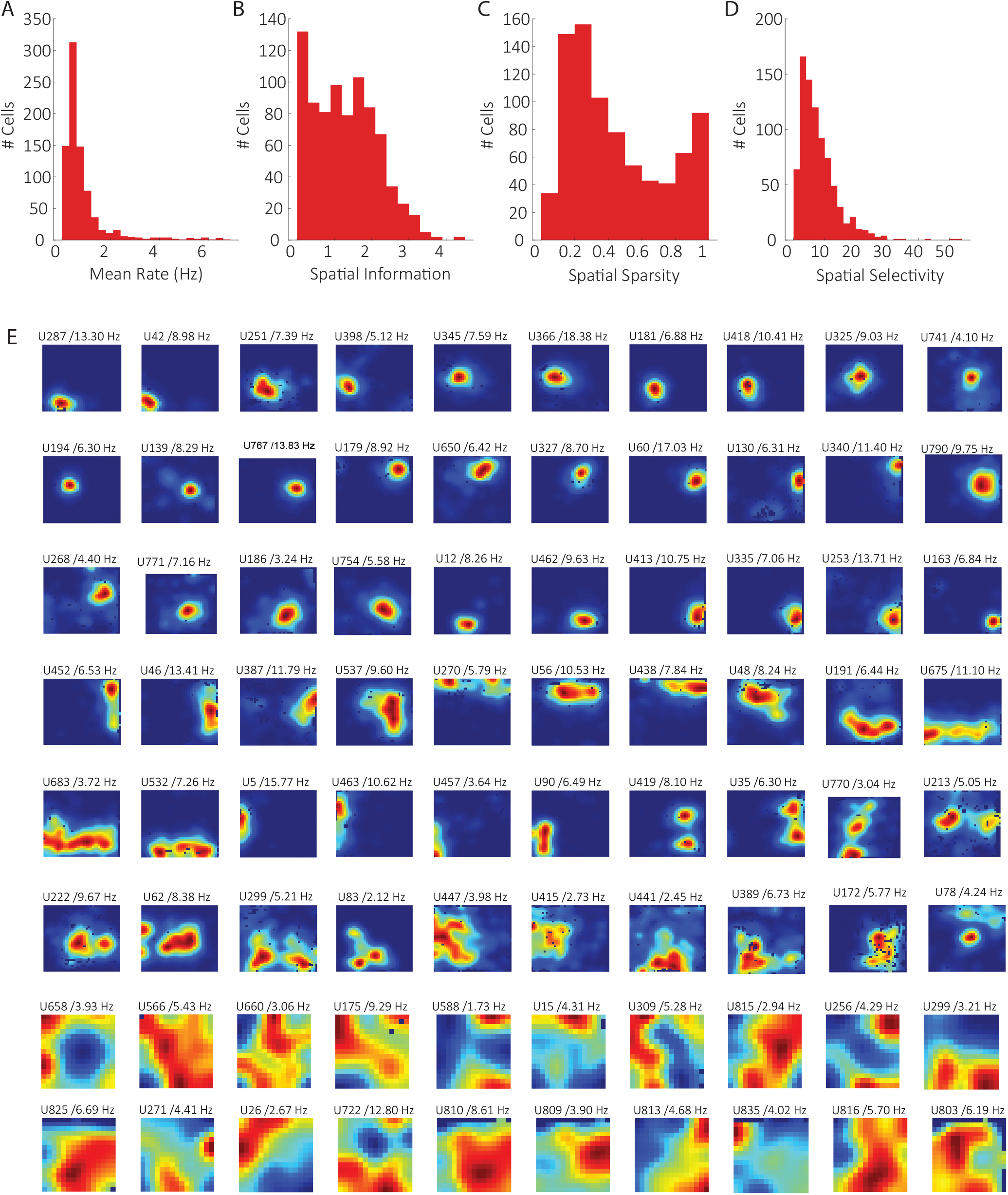
Properties of recorded neurons in the dataset. **A**. Mean firing rate (in Hertz) for each recorded neuron. **B**. Spatial information score for each recorded neuron. **C**. Spatial sparsity score for each recorded neuron. **D**. Spatial selectivity for each recorded neuron. **E**. Spatial ratemaps from example cells recorded across all animals. The numbers on top of each plot indicate the unit number (U) and the peak firing rate of the spatial ratemap in Hertz. Note that our dataset is made up of place cells with small firing fields, place cells that primarily fire along borders, and broadly spatially tuned neurons.

**Supplemental Figure 3:**
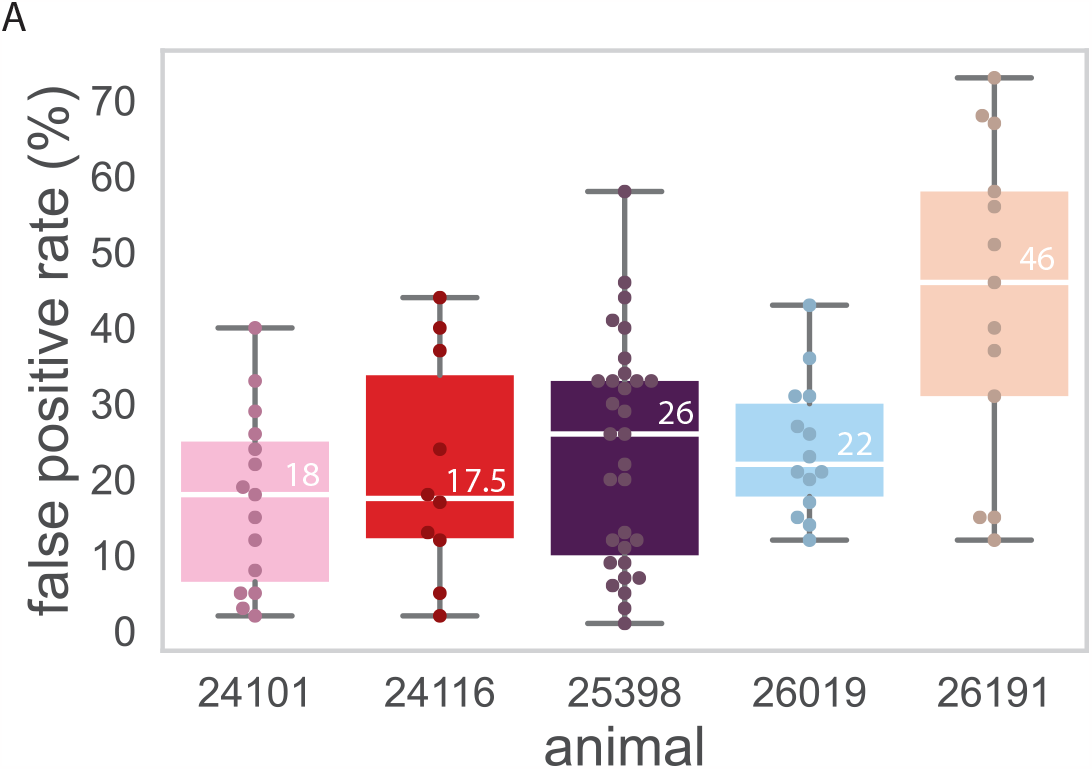
False positive rates vary across animals. **A**. Distribution of false positive rates (%) across animals, each labeled by their unique I.D. number. The central mark indicates the median, and the bottom and top edges of the box indicate the 25th and 75th percentiles, respectively. The whiskers extend to the most extreme data points not considered outliers. All data points (gray dots) are shown overlaid.

**Supplemental Figure 4:**
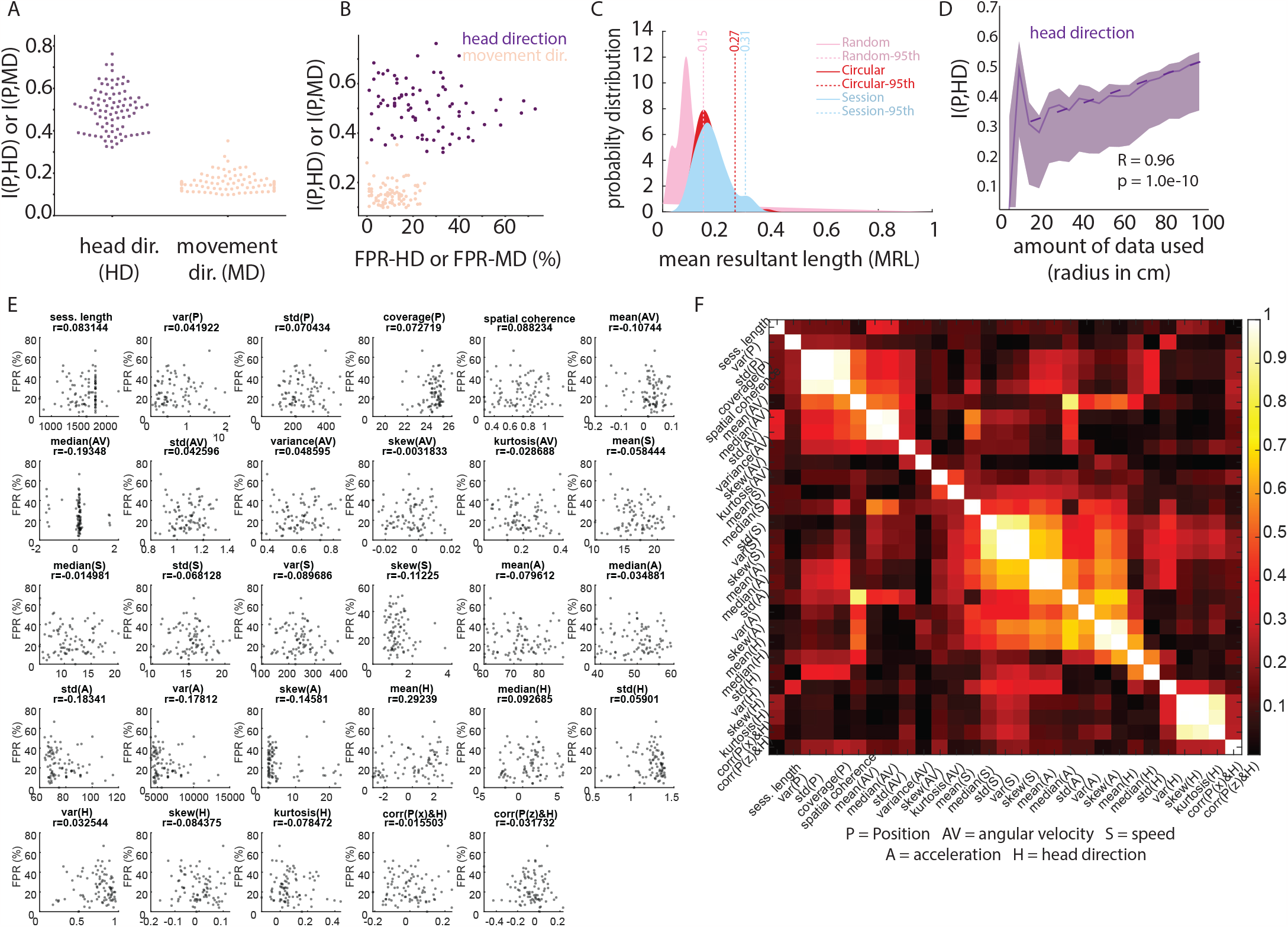
Exploring variables related to variations across false positive rates. **A**. Median mutual information values (calculated for all 86 open field behavioral sessions) between position and head direction I(P,HD) (purple) and mutual information between position and movement direction I(P,MD) (orange) arranged in a swarm plot. **B**. False positive rate (%) calculated by deriving egocentric bearing relative to the head direction (FPR-HD) or movement direction (FPR-MD) vs. mutual information between the animal’s position and head direction, I(P,HD) or movement direction, I(P,MD). Each datapoint is a session, and each session is colored by the rat. Note that there is no strong linear correlation between these two variables, meaning that one alone does not predict the other. **C**. Distribution of mean resultant lengths of RH-model fit tuning curves for a single cell, when applying three different null hypotheses (and thus permutation tests). In all cases, the spike train is held constant relative to the rat’s head direction. The distribution is built in pink when head direction estimates are randomly shuffled. The dashed yellow line is the 95th percentile of the null distribution (calculated non-parametrically). The results of circularly permuting the head direction indices are in red, and in blue, the head direction estimates from another session are used to construct the null. **D**. Mutual information between position and head direction I(P,HD) (purple) is calculated from data taken from larger and larger circular portions (quantified by radius) of the 2D recording arena, starting in the center. Aside from expected increases in mutual information between radius = 0 and radius = 20 due to extremely small amounts of data, I(P,HD) increase strongly when data from the borders of the environment are incorporated into the measure (e.g., when the radius increases to reach the size of the recording environment). **E**. Twenty-nine variables were calculated using the experimentally recorded behavior from the 86 open field sessions. Most of the variables were statistics of head direction (H), angular velocity (AV), speed (S), and acceleration (A). Other variables included a metric of the animal’s coverage (BNT), session length, and spatial coherence. For each session, we plotted each of the 29 variables as a function of the false positive rate for that session (HD-CIRC). Above each plot is the linear correlation between the FPR and given variable (Pearsons r). No significant linear correlations were found between FPR and any single behavioral variable. **F**. Correlation matrix of the 29 calculated variables to show that many co-vary with each other (color axis = correlation coefficient).

**Supplemental Figure 5:**
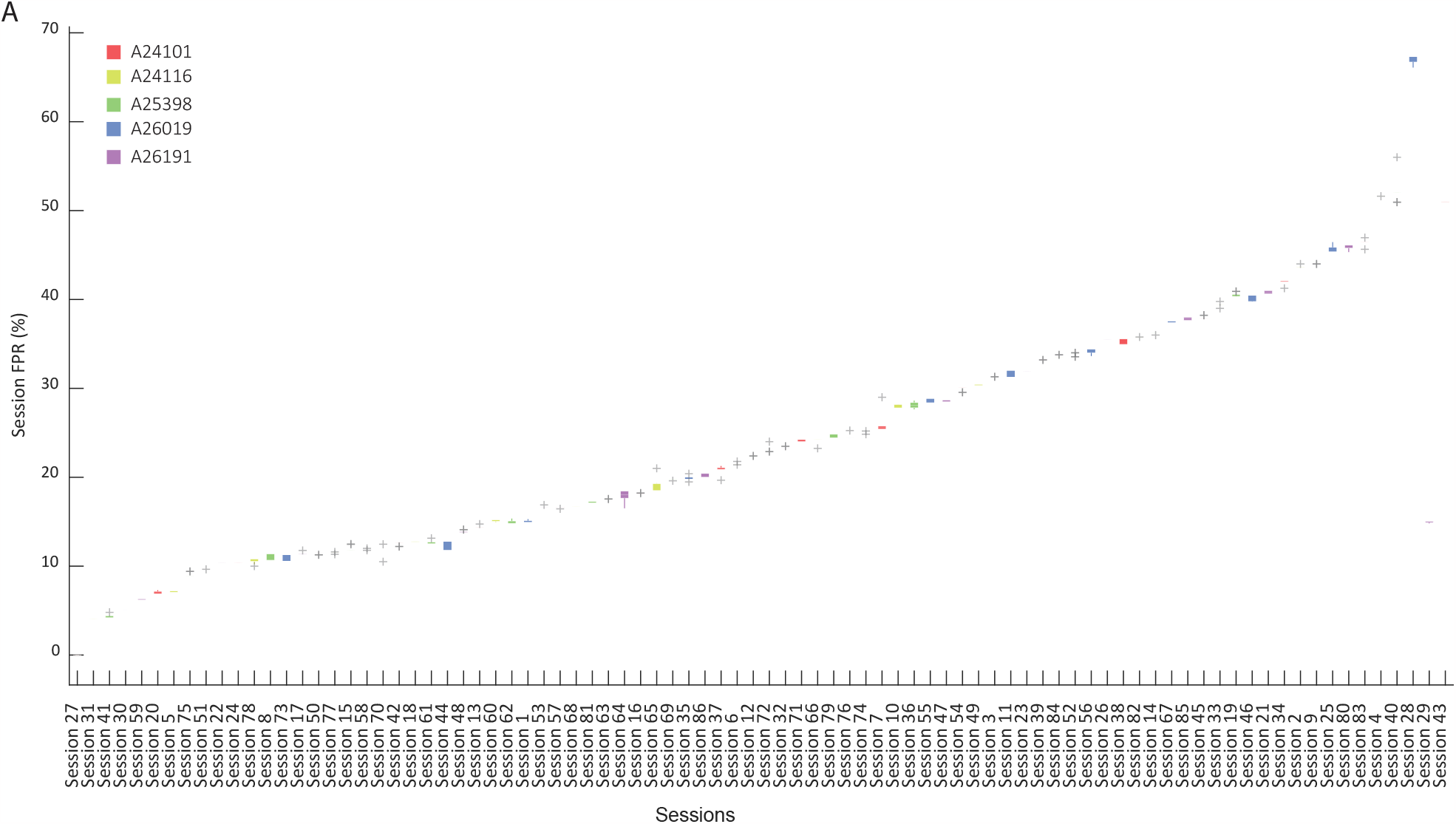
False positive rate for single sessions is consistent across multiple runs. **A**. Boxplots of the distributions of false positive rates for each session, sorted from sessions with the lowest false positive rates, to those with the highest false positive rates. Boxplots are colored by the animal that the session was recorded from.

**Supplemental Figure 6:**
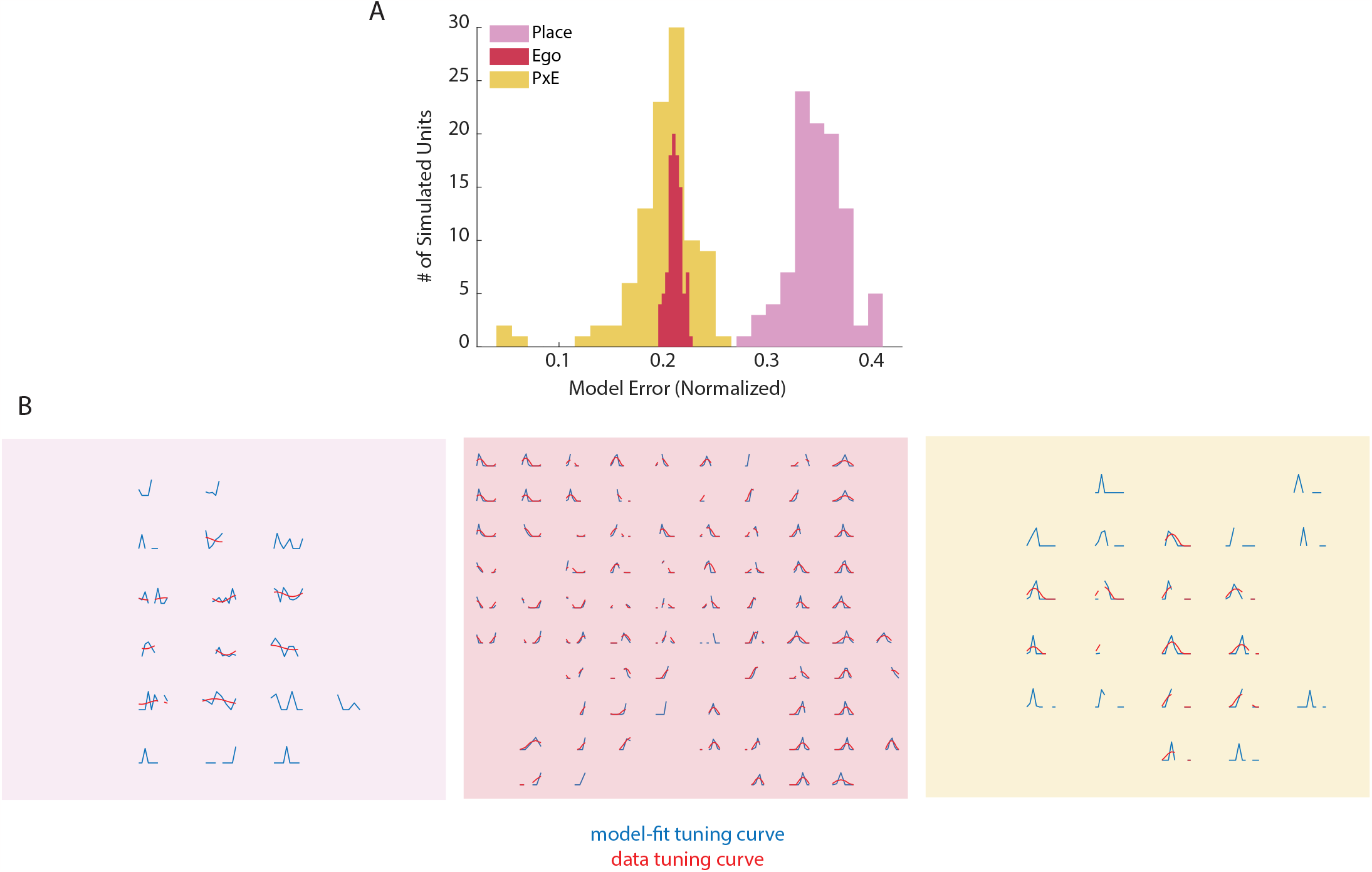
Model error is much higher for simulated place cells than egocentric cells. **A**. Histograms showing the model error (MSE) for 100 simulated cells from session #1. The simulated cells were either place cells (yellow), egocentric cells (orange), or conjunctive place + egocentric cells (blue). We found a statistically significant difference between the three distributions (p¡0.05 for all combinations; Wilcoxon Rank-Sum Test). **B**. Examples of the tuning curve constructed from the data (blue) vs. the model-fit cosine curve (red) for each spatial bin where the simulated cells fired. From left to right we show an example simulated place cell, an egocentric cell, and a conjunctive egocentric + place cell.

**Supplemental Figure 7:**
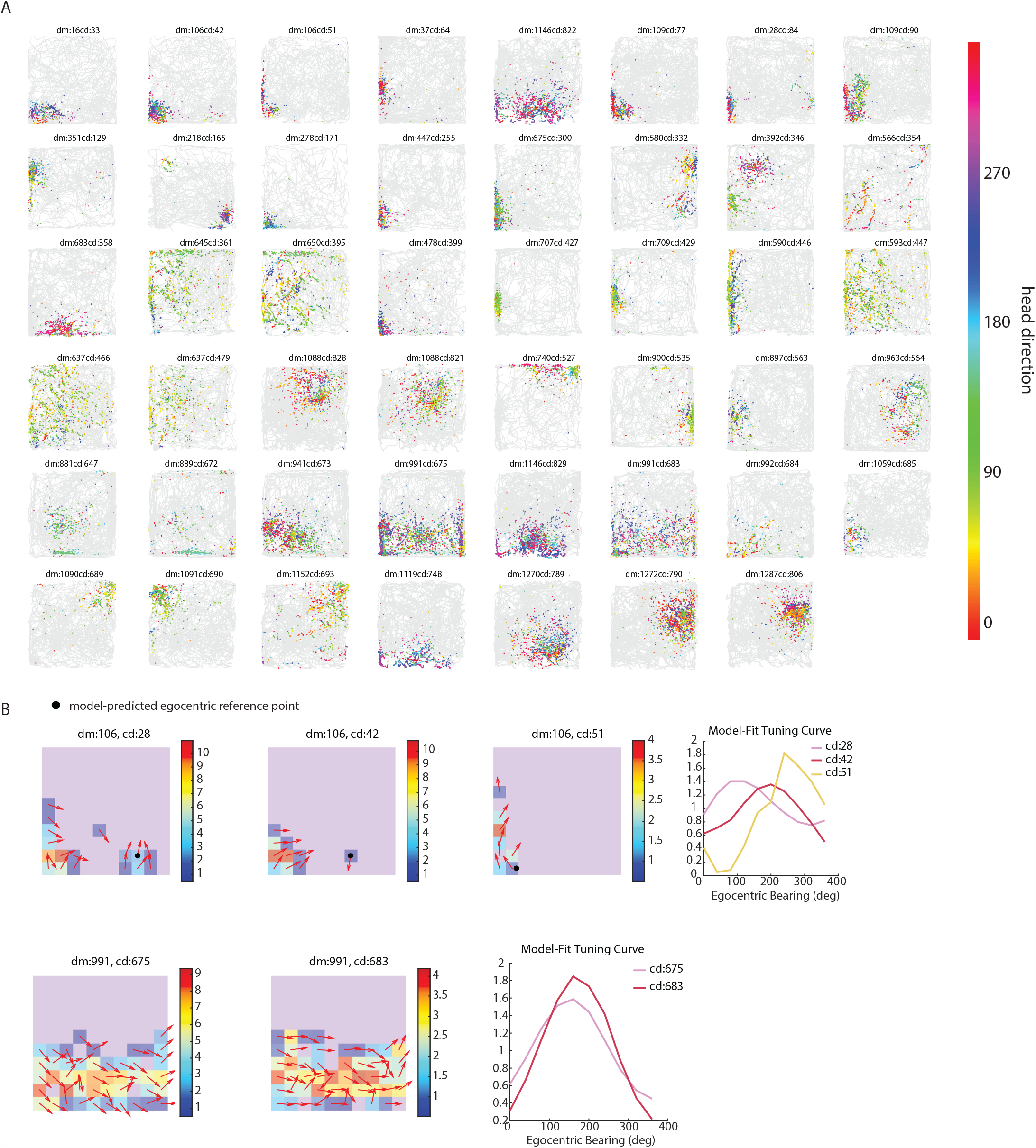
Egocentric-modulated neurons according to copycat-neuron tests. **A**. Example head-direction colored spike plots of recorded CA1 neurons that are significantly more EB-modulated than their respective copycat distributions. Plotted above each neuron is the unique identifier of neurons tracked across sessions (‘dm’) and another identifier that acts as a count (‘cm’). **B**. Within the group of neurons showcased in section **A**, only two consistently responded to egocentric bearing in multiple sessions: Neurons numbered 106 and 991. These two passed the strictest selection criteria in at least two separate sessions. For each neuron, the left-side diagrams display a simplified map of their firing rates across different locations, with colors indicating the intensity of the firing rate. The red arrows are the preferred head direction (according to the RH model) in each spatial bin. The black dot delineates the optimal egocentric reference point predicted by the RH-model. The right plots show the predicted egocentric tuning curves for each recorded instance of the cell.

